# Visual feature tuning properties of short-latency stimulus-driven ocular position drift responses during gaze fixation

**DOI:** 10.1101/2023.09.25.559257

**Authors:** Fatemeh Khademi, Tong Zhang, Matthias P. Baumann, Tatiana Malevich, Yue Yu, Ziad M. Hafed

## Abstract

Ocular position drifts during gaze fixation are generally considered to be random walks. However, we recently identified a short-latency ocular position drift response, of approximately 1 min arc amplitude, that is triggered within <100 ms by visual onsets. This systematic eye movement response is feature-tuned and seems to be coordinated with a simultaneous resetting of the saccadic system by visual stimuli. However, much remains to be learned about the drift response, especially for designing better-informed neurophysiological experiments unraveling its mechanistic substrates. Here we systematically tested multiple new feature tuning properties of drift responses. Using highly precise eye tracking in three male rhesus macaque monkeys, we found that drift responses still occur for tiny foveal visual stimuli. Moreover, the responses exhibit size tuning, scaling their amplitude as a function of stimulus size, and they also possess a monotonically increasing contrast sensitivity curve. Importantly, short-latency drift responses still occur for small peripheral visual targets, which additionally introduce spatially-directed modulations in drift trajectories towards the appearing peripheral stimuli. Drift responses also remain predominantly upward even for stimuli exclusively located in the lower visual field, and even when starting gaze position is upward. When we checked the timing of drift responses, we found that it was better synchronized to stimulus-induced saccadic inhibition timing than to stimulus onset. These results, along with a suppression of drift response amplitudes by peri-stimulus saccades, suggest that drift responses reflect the rapid impacts of short-latency and feature-tuned visual neural activity on final oculomotor control circuitry in the brain.

**Significance:** During gaze fixation, the eye drifts slowly in between microsaccades. While eye position drifts are generally considered to be random eye movements, we recently found that they are modulated with very short latencies by some stimulus onsets. Here we characterized the feature-tuning properties of such stimulus-driven drift responses. Our results demonstrate that drift eye movements are not random, and that visual stimuli can impact them in a manner similar to how such stimuli impact microsaccades.

## Introduction

The eye is never completely still during gaze fixation (Barlow, 1952; Steinman et al., 1967; Steinman et al., 1973), resulting in subtle, but continuous, alterations of the retinal image streams entering the visual system. Two primary components of fixational eye movements are microsaccades and slow ocular position drifts (Fig. 1A). While the neural control of microsaccades is relatively well established (Krauzlis et al., 2017; Hafed et al., 2021a), that of ocular position drifts is less understood. Moreover, the ways with which external sensory transients interact with these two types of eye movements are not fully investigated.

**Figure 1.**
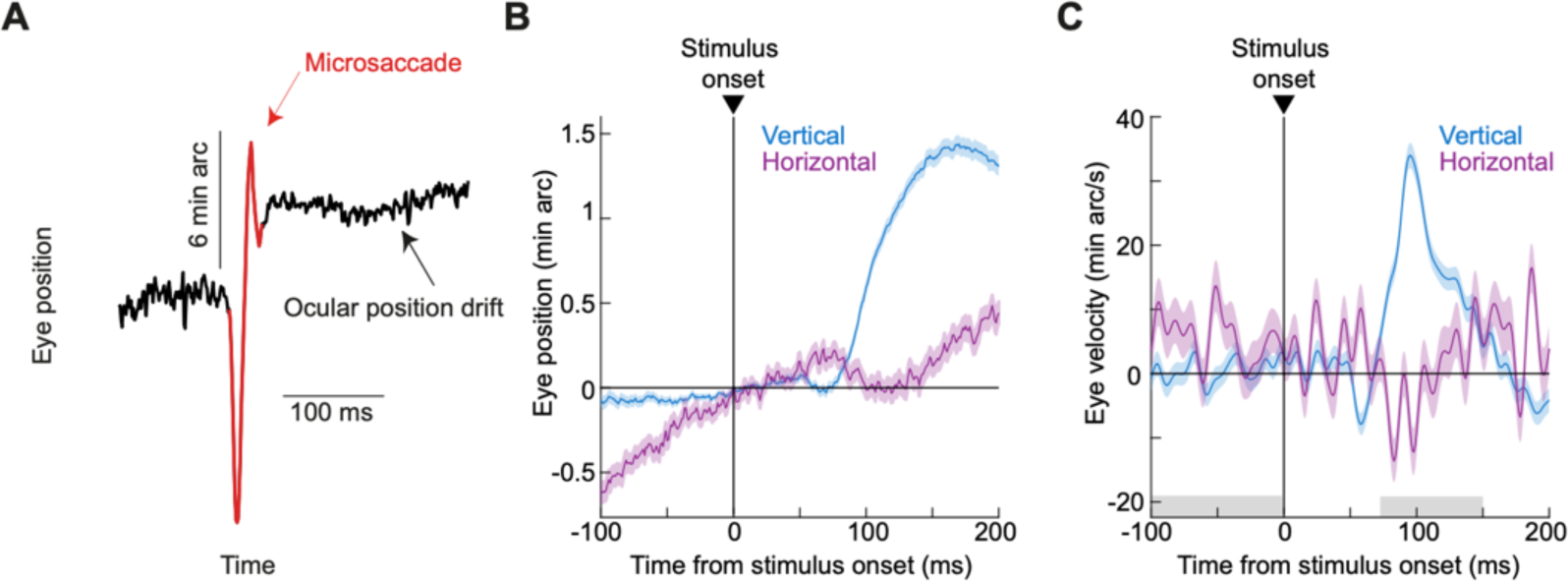
Stimulus-driven ocular position drift responses. **(A)** Accurate gaze fixation is characterized by two prominent features: (1) microsaccades occur occasionally to re-align the line of sight (red); and (2) the eye drifts continuously with slow speeds in between saccades and microsaccades (black). **(B)** We recently found (Malevich et al., 2020) that large stimulus onsets result in a short-latency change in ocular position drift statistics, primarily marked by a small upward deviation in eye position (although an earlier, even smaller, downward movement component jumpstarts the whole response sequence). The figure shows average horizontal and vertical eye positions (surrounded by SEM ranges; n = 882 trials) from an example condition and an example monkey (A) from Experiment 1 of the current study, replicating (Malevich et al., 2020). Positive deflections in each curve indicate rightward and upward eye position deviations, respectively, and the data across trials were first aligned to eye position at time zero before averaging (Malevich et al., 2020) (Materials and Methods). As can be seen, the monkey exhibited rightward pre-stimulus drifts; after stimulus onset, there was a predominantly upward drift response, which was accompanied by a small leftward component to it. The upward drift response was also preceded by a much smaller and shorter-lived downward eye position deviation, although we primarily focus here on the overall upward nature of the whole response sequence. **(C)** Horizontal and vertical eye velocity curves (surrounded by SEM ranges) from the same trials as in **B**. The stimulus-driven drift response was predominantly upward. Shaded regions on the x-axis indicate our measurement intervals of baseline (pre-stimulus) and post-stimulus eye velocities, for use in our summary statistics in the remainder of this article.

For microsaccades, visual transients in the environment rapidly reset the oculomotor rhythm, causing microsaccadic inhibition (Engbert and Kliegl, 2003; Hafed et al., 2021b; Buonocore and Hafed, 2023), and giving rise to important implications on subsequent perceptual performance and visual neural sensitivity (Hafed et al., 2015). Moreover, such inhibition is feature-tuned, altering its time course and strength as a function of the appearing visual patterns (Khademi et al., 2023). This likely reflects the tuning properties of visually-sensitive neurons mediating microsaccadic inhibition (Buonocore and Hafed, 2023).

For drifts, we recently found that certain visual stimuli robustly trigger a short-latency change in drift statistics, which we refer to here as the drift response (Malevich et al., 2020). This response is characterized by a small predominantly upward displacement, superseding the ongoing drift direction, and being much slower than even the slowest microsaccades. For example, in Fig. 1B, aligning all eye position epochs at the time of stimulus onset reveals a predominantly rightward drift trajectory prior to stimulus onset; this rightward drift was momentarily transformed into a predominantly upward drift pulse within less than 100 ms after stimulus onset, with an even smaller downward component just prior to that (Fig. 1B, C) (Malevich et al., 2020).

Our previous work revealed that the drift response occurred when we presented relatively large stimuli (Malevich et al., 2020). We also found that this drift response, much like saccadic inhibition (Khademi et al., 2023), is feature-tuned. Specifically, it was stronger for low spatial frequency patterns, as well as for certain grating orientations (Malevich et al., 2020). However, understanding the full mechanisms underlying the drift response requires much deeper characterization of this response’s functional properties. For example, might such a drift response still occur for small visual stimuli, just like microsaccades can be affected by small eccentric targets (Hafed and Clark, 2002; Engbert and Kliegl, 2003)? And, would the predominantly upward nature of the drift response change if we only presented lower visual field stimuli rather than stimuli spanning both sides of the retinotopic horizon?

Here we answered these, and other, questions, and we laid down a rich foundation for testing the neurophysiological underpinnings of not only the drift response, but also of the coordination between multiple types of fixational and targeting eye movements with external sensory events. We first found that the drift response is size-tuned, and can still happen for tiny, foveal visual stimuli. We also characterized the contrast sensitivity of the drift response, as well as its modulation by small peripheral visual targets. Interestingly, and unlike our expectation (Malevich et al., 2020) that the drift response might reflect the preference of the superior colliculus (SC) for the upper visual field (Hafed and Chen, 2016; Fracasso et al., 2023), we found that the drift response is still predominantly upward even for stimuli below the horizon. Finally, we characterized the temporal coordination between microsaccades and the drift response, as well as the alteration of the drift response magnitude by the occurrence of peri-stimulus microsaccades, mimicking the classic phenomenon of saccadic suppression (Zuber and Stark, 1966; Beeler, 1967; Hafed and Krauzlis, 2010; Idrees et al., 2020).

Our results demonstrate that the “lens” through which the oculomotor system processes visual scenes may be similar for dictating the visual feature tuning properties of both saccadic inhibition (Khademi et al., 2023) and drift responses, and that these two ubiquitous eye movement phenomena likely arise from a common underlying source.

## Materials and methods

### Experimental animals and ethical approvals

We collected data from three adult, male rhesus macaque monkeys (*macaca mulatta*), referred to here as A, F, and M, respectively. The monkeys were aged 7-14 years, and they weighed 9.5-12.5 kg. All experiments were approved by ethics committees at the regional governmental offices of the city of Tübingen.

### Laboratory setup and animal procedures

Some experiments involved analysis of ocular position drifts from our recent study, which only focused on saccades (Khademi et al., 2023). Other experiments were run specifically for the purposes of the current study, but in the same experimental setups as in (Khademi et al., 2023). The reader is referred to our recent publication for details on our laboratory equipment (Khademi et al., 2023). Briefly, we used precise eye tracking, using the scleral search coil technique (Robinson, 1963; Fuchs and Robinson, 1966; Judge et al., 1980), and a real-time experimental control system based on PLDAPS (Eastman and Huk, 2012) and the Psychophysics Toolbox (Brainard, 1997; Pelli, 1997; Kleiner et al., 2007). The monkeys had their heads stabilized during the experiments, and they watched stimuli on a computer-controlled display in front of them. The display size was spanning approximately 31 deg horizontally and 23 deg vertically, and the experimental room was otherwise dark.

### Experimental procedures

The experiments all involved gaze fixation, and we analyzed fixational eye movements. The experimental procedures were described in detail recently (Khademi et al., 2023). In brief, the monkeys fixated a small, stationary fixation spot presented over a gray background (of luminance 26.11 or 36.5 cd/m^2^). At a random time during fixation, a single-frame flash (∼12 or ∼7-8 ms) was presented. Across trials and experiments, the flash could have different feature properties (for example, full-screen flash or small, localized target, and so on). In what follows, we describe the experiment-specific details, explaining what image features the brief flashes had in the different experiments.

#### Experiment 1: Size tuning

This experiment was the same as that used recently (Khademi et al., 2023). In that study, we analyzed the saccades that took place around stimulus onset. In the current study, we analyzed ocular position drifts (in saccade-free epochs), as well as saccade-drift interactions, as we describe in more detail below.

The stimulus flash in this experiment consisted of a black circle of different radii across trials. The range of sizes tested included stimuli approximately as small as the fixation spot (0.09 deg radius), stimuli approximately as large as the entire display (9.12 deg radius), and stimuli with sizes in between these two extremes. Moreover, the numbers of trials collected were the same as those reported in (Khademi et al., 2023).

For the numbers of trials that were analyzed, these depended on whether we picked drift response trials (saccade-free) or trials with peri-stimulus microsaccades (see Data Analysis below for details). For example, as we describe in more detail below, for some analyses, we only considered trials in which there were no microsaccades in the interval from -100 ms to 200 ms relative to stimulus onset, and in some other analyses, we considered trials with microsaccades happening in the final 100 ms before stimulus onset, and so on. That is why we document the specific numbers of trials included in the analyses of each figure shown in Results separately.

#### Experiment 2: Contrast sensitivity with full-screen stimuli

This experiment was again the same as that used recently (Khademi et al., 2023). Briefly, the stimulus onset could be a full-screen flash having one of five different Weber contrasts (5%, 10%, 20%, 40%, or 80%). Once again, we analyzed saccade-free drift response trials as well as trials having saccades within specific time intervals relative to stimulus onset (see Data Analysis below for more details). For each analysis, the numbers of trials included are documented individually in Results. Drift-only (saccade-free) trials were not analyzed previously in (Khademi et al., 2023).

#### Experiment 3: Upper and lower visual field stimuli

This experiment was collected specifically for this study (as well as related ongoing neurophysiological experiments). The general trial sequence was the same as that in the above two experiments. Specifically, the monkeys fixated a central spot. After a random time, one of five different events took place, depending on the trial type. The first trial type was just a sham condition: no stimulus display update occurred at all, but we just used the sham event in the data file to study baseline drift trajectories and compare them to trajectories with a real stimulus. The second trial type had the stimulus being a 1 deg x 1 deg black square that was flashed for a single display frame. The location of the flash was somewhere in the periphery relative to the central fixation spot (approximately 3.5-11 deg), but this location was constant within a given session. This location was typically dictated by the locations of receptive fields of neurons that we were recording simultaneously for other purposes, since this task was typically run while we recorded SC and/or primary visual cortex activity. The third trial type was a 100% black full-screen flash (again with a duration of a single frame). Here, the stimulus was basically similar to the stimuli used in Experiment 2 above. And, finally, the fourth and fifth trial types were half-screen flashes. Specifically, we split the screen in half along the vertical dimension. In one condition, the flash was only in the upper half of the screen (above the midline defined by the vertical position of the fixation spot), and in another condition, the flash was only in the lower half of the screen.

We typically ran this task in daily blocks of approximately 100-500 trials per session, and we collected a total of 7524, 7521, and 7495 trials in monkeys A, F, and M, respectively. This resulted in 72-1208 trials per condition per animal for the saccade-free drift response analyses (like in Fig. 1B, C).

#### Experiment 4: Small, localized stimuli across different visual field directions

Because the locations of the small stimuli used in Experiment 3 were dictated by other experimental constraints (such as receptive field locations), we ran an additional experiment in which we sampled eccentric locations more evenly. Specifically, the experiment consisted of the transient flash being a 1 deg x 1 deg black square at a 7.9 deg eccentricity from the display center. The square could appear in one of 8 equally spaced directions, thus covering both right and left as well as up and down visual field locations. The flash location was randomly interleaved across trials.

We typically ran this task in daily blocks of 310-900 trials per session, and we collected a total of 5961, 4357, and 6048 trials in monkeys A, F, and M, respectively. This resulted in 65-383 analyzed trials per location per animal for the basic saccade-free drift response analyses. We typically pooled multiple locations for a given analysis, as we describe below, in order to increase statistical confidence in the results. Once again, all numbers of trials are documented in appropriate sections of Results.

#### Experiment 5: Gaze position

This task was the same as that in Experiment 2 above, with only one difference. Across sessions, the fixation spot could be at 4 deg to the right, left, up, and down relative to the display center. This task, therefore, allowed us to test whether the drift response (Fig. 1B, C) was substantially different if the starting gaze position of the eye was different.

We ran 4 sessions of this task in monkey A, collecting a total of 2206 trials. This resulted in 500-602 analyzed trials per eye position for the basic saccade-free drift response analyses.

### Data analysis

All saccades were analyzed as described recently (Khademi et al., 2023). Briefly, we detected saccades of all sizes using our established methods (Chen and Hafed, 2013; Bellet et al., 2019), and we included all detected saccades that took place around stimulus onset. This allowed us to estimate saccadic inhibition latency using the L_50_ parameter (Reingold and Stampe, 2002, 2004; Rolfs et al., 2008; Khademi et al., 2023). Simply put, this parameter describes when the saccade rate curve drops by 50% of the dynamic range between pre-stimulus (baseline) saccade rate and the minimum saccade rate during saccadic inhibition. The reader is referred to our detailed description of this parameter in (Khademi et al., 2023). We estimated saccade rate using the method described in (Khademi et al., 2023): briefly, we calculated saccade onset likelihood within 50 ms moving windows that were stepped in time by 1 ms steps, and we did this on a per-trial basis; across-trial average rates were then obtained in order to calculate L_50_ from the global saccade rate. While we acknowledge that there might be other means to estimate the latency of saccadic inhibition (Bompas et al., 2023), we used L_50_ because of its consistent use in other studies (Reingold and Stampe, 2002, 2004; Rolfs et al., 2008; Khademi et al., 2023), and also because it does a good job in capturing the drop in saccade likelihood across conditions (see, for example, Fig. 7 later in Results).

To visualize drift responses, we averaged the horizontal and vertical eye position traces of a given animal and condition across trial repetitions. Before such averaging, we realigned each trace to the position of the eye at the time of stimulus onset (Malevich et al., 2020). This allowed us to isolate visualization of the drift statistics despite variations in absolute eye position at the time of stimulus onset, due to continuous fixational eye movements. We also visualized drift responses by plotting vertical eye velocity traces (e.g. Fig. 1C). We obtained these traces using a smooth differentiating filter (Chen and Hafed, 2013; Malevich et al., 2020) applied to vertical eye position on a trial-by-trial basis. We then averaged the individual trial velocity traces.

For all analyses characterizing the drift response, we only picked trials without any saccades in the interval from -100 ms to 200 ms relative to stimulus onset. This was done for two reasons: to avoid masking the slow drift responses by large velocity pulses associated with saccades, and to avoid potential peri-saccadic modulations in the drift response strength. In some analyses, we specifically wanted to study such peri-saccadic modulations, as well as drift-saccade interactions in general. In that case, we replaced all velocity samples that were part of a saccade with not-a-number (NaN) labels before averaging the eye velocity traces across trials.

For summary statistics, we estimated the size of the drift response by calculating average vertical eye velocity in a post-stimulus response interval (70-150 ms; second gray interval on the x-axis in Fig. 1C) and subtracting from it the baseline vertical eye velocity in a pre-stimulus interval (first gray interval on the x-axis in Fig. 1C). We did this on a trial-by-trial basis, and we then averaged the difference measures across trials for population statistics. Note that this velocity difference measure could quantitatively be negative, especially in the cases with weak or non-existent drift responses (Malevich et al., 2020). Note also that we picked the post-stimulus response interval (70-150 ms) by inspecting drift responses across many different trials, conditions, and animals. While this interval was fixed for all analyses, it was long enough to avoid biasing our results in the cases in which the drift response was rendered a bit earlier or a bit later by specific visual feature dimensions.

For analyzing the impacts of peri-stimulus saccades on the drift response, we calculated the response strength measure just described above but now only for trials in which saccade onsets occurred within a specific time window relative to stimulus onset. This time window was defined by the purposes of the specific analysis (see Results).

Finally, for analyzing effects of localized flash locations on drift responses, we sometimes also measured eye position rather than eye velocity. In this case, we grouped trials according to whether a flash was in the right or left visual field (independent of its vertical position), and we took the difference in eye position (after aligning all traces at time zero like above) between the two groups of trials in a given post-stimulus interval. Similarly, we also grouped trials according to whether a flash was in the upper or lower visual field (independent of its horizontal position), and we took the difference in eye position between the two groups of trials (again, after all traces were aligned at the time of stimulus onset, like described above). Using eye position instead of eye velocity in these particular analyses allowed us to directly test whether there were spatially-directed modulations in drift statistics that were caused by eccentric stimulus onsets (see Results), similar to how eccentric stimulus onsets can bias microsaccade directions (Hafed and Clark, 2002; Engbert and Kliegl, 2003).

#### Experimental design and statistical analyses

We always replicated all of our results in three monkeys (except for Experiment 5; see justification below). Moreover, within each animal, we typically had hundreds to thousands of trial repetitions per condition (see, for example, Fig. 1). This increased our confidence in our population measures. Our choice of trial numbers to collect was guided by calculating power estimates before and during the experimental phases of the study. We also randomly interleaved all conditions in a given experiment, except when we were constrained by the experimental setup. For example, in Experiment 3, the location of the small, localized flashes was constant within a given session, and this was dictated by other factors external to the study (like receptive field locations). However, given the reflexive nature of our drift responses (see Results and Discussion), this should not have affected our interpretations in any substantial manner. More importantly, we also designed Experiment 4 with randomly interleaved target locations exactly to compensate for the non-random nature of localized flash locations in Experiment 3.

For Experiment 5, we only ran it in one monkey. However, the results were virtually identical, in a qualitative sense, to everything else that we had tested with the other two animals in other experiments. As a result, we decided that our conclusions from this experiment were already convincing. Similarly, we blocked gaze position in this experiment, meaning that we tested each gaze position condition in a block of contiguous trials (as opposed to randomly changing gaze position from trial to trial). Again, this provided a stronger support for our conclusions that the drift response remains to be predominantly upward independent of gaze position (see Results).

All statistical tests and outcomes, as well as trial repetition counts, are detailed in Results. We also performed statistical tests for each animal separately.

## Results

We recently found that ocular position drifts can be quite sensitive to visual stimulus onsets, exhibiting short-latency, brief responses (Fig. 1) (Malevich et al., 2020). Here, we performed extensive additional experiments characterizing the feature tuning properties of such stimulus-driven drift responses.

We used three rhesus macaque monkeys as our experimental subjects, and we did so for at least four reasons. First, we employed highly precise eye tracking in these animals, using the scleral search coil technique (Robinson, 1963; Fuchs and Robinson, 1966; Judge et al., 1980), to increase our confidence in the measurements. Commercial video-based eye trackers commonly used with human subjects would make measuring these tiny drift responses very challenging (Wyatt, 2010; Kimmel et al., 2012; Chen and Hafed, 2013; Choe et al., 2016; Malevich et al., 2020). Second, we could collect several experimental sessions per animal per condition, resulting in many trial repetitions and statistically robust results across all of our experimental conditions (Materials and Methods). Third, these animals were already used in our characterization of the closely related phenomenon of saccadic inhibition (Khademi et al., 2023), and we often used the very same data for characterizing drift responses here. Fourth, and most importantly, these animals are part of the ongoing efforts in our laboratory to explore the neurophysiological underpinnings of drift responses, which we hope to document in the near future.

### The drift response exhibits size tuning

In our first experiment, we asked whether the ocular position drift response is parametrically tuned to the size of the appearing visual stimulus. In our initial characterization of the drift response (Malevich et al., 2020), we mostly used large visual stimuli (full or half of our experimental stimulus displays). This raises the question of how small the visual target needs to be for the drift response to disappear. We instructed our monkeys to maintain fixation on a central fixation spot, and we presented a brief flash of a black circle centered on the fixation spot (Materials and Methods). The flash could be approximately as small as the fixation spot or as large as the entire display, with intermediate radii in between, and we analyzed data from the same experiments in which we recently characterized saccadic inhibition as a function of stimulus size (Khademi et al., 2023). The difference in the current study is that we specifically focused here on trials in which there were no microsaccades occurring within the interval between -100 ms and 200 ms from stimulus onset (Materials and Methods; also see later for our separate analyses investigating interactions between microsaccades and the drift response).

The smallest foveal visual stimulus could still evoke a clear drift response. Figure 2A, B (yellow) shows average horizontal (Fig. 2A) and vertical (Fig. 2B) eye position from monkey A when the smallest visual flash occurred. In each panel, we always aligned all eye position traces across trials to the eye position at time zero (stimulus onset), in order to isolate the impact of the stimulus event on drift statistics (despite variable eye positions during gaze fixation; Materials and Methods) (Malevich et al., 2020). As can be seen, this monkey had a systematic rightward drift trajectory before stimulus onset (Fig. 2A, yellow); that is, the horizontal eye position curve in Fig. 2A was steadily shifting upward in the plot (meaning a rightward displacement) during the pre-stimulus interval; the vertical eye position curve in Fig. 2B was more-or-less steady. After stimulus onset, Fig. 2B shows that there was still a small upward drift response that occurred (not unlike that seen in Fig. 1B, C), despite the vanishingly small stimulus size relative to the size of the fixation spot. Such a small upward drift response was also clearly visible in monkey F (Fig. 2D, E, yellow curves), even though this monkey had a different pre-stimulus drift trajectory (which was now predominantly leftward and downward). In monkey M, the smallest visual stimulus barely modified the ongoing drift statistics (Fig. 2G, H, yellow curves), but this monkey also had the fastest pre-stimulus drift speeds from among all three animals (compare the rates of change in eye positions during the pre-stimulus epochs across all panels). This faster baseline drift speed might have masked any potential impacts of the smallest stimulus size on drift eye movements in this monkey. Nonetheless, and as we describe next, drift responses were still clearly visible in this animal for the slightly larger stimulus radii of only 0.18 or 0.36 deg. Thus, in all three animals, even the smallest, foveal stimuli could still evoke a reliable, predominantly upward, drift response.

**Figure 2.**
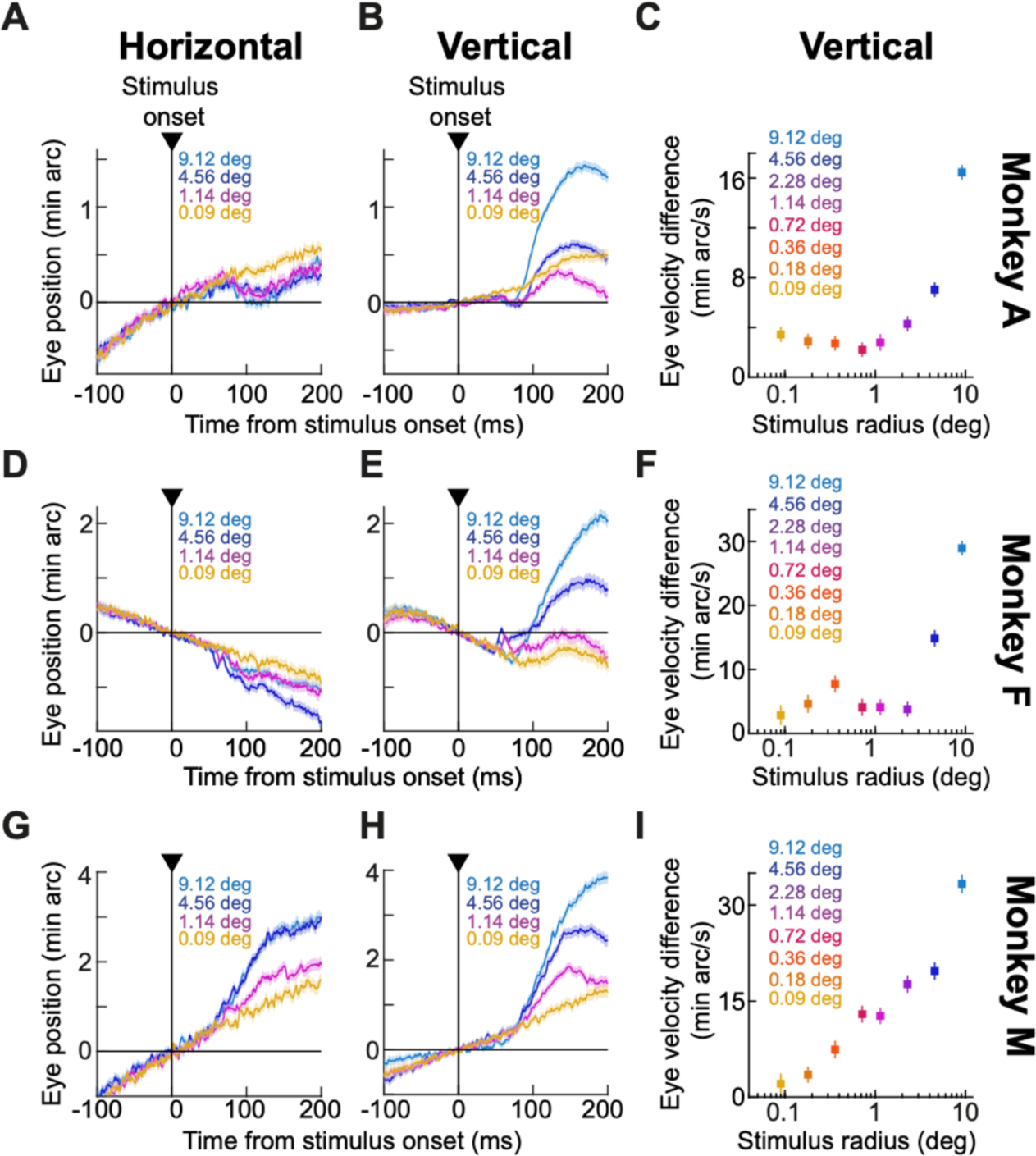
Size tuning of ocular position drift responses. **(A)** Average horizontal eye position from monkey A for four example stimulus sizes (0.09 deg, 1.14 deg, 4.56 deg, and 9.12 deg). Error bars denote SEM, and the numbers of trials were 827, 804, 927, and 882 for the four stimulus sizes, respectively. Upward deflections in the plot denote rightward eye position deflections. **(B)** Average vertical eye position from the same trials as in **A**; error bars again denote SEM, and upward deflections denote upward eye position deflections. A clear dependence of the ocular position drift response on stimulus size can be seen. Note also how the smallest tested stimulus (0.09 deg) still caused a vertical drift response, but its initial smaller downward component was missing. **(C)** Our measure of the drift response magnitude (average baseline-corrected vertical eye velocity in the interval 70-150 ms after stimulus onset; Fig. 1C; Materials and Methods) for all tested stimulus sizes in monkey A (n = 827, 729, 872, 868, 804, 885, 927, and 882 trials from the smallest to the largest stimulus size). Error bars denote SEM. **(D-F)** Similar results for monkey F (n = 223, 219, 235, 266, 308, 339, 350, and 399 trials from the smallest to the largest stimulus size). Note how this monkey also showed small transient oscillations in both horizontal and vertical eye positions at the very initial phases of the drift response. **(G-I)** Similar results for monkey M (n = 327, 369, 397, 423, 456, 420, 416, and 405 trials from the smallest to the largest stimulus size). In all monkeys, the drift response was size-dependent, and it increased monotonically with sizes beyond 1-2 deg.

The drift response not only occurred for small, foveal stimuli, but its magnitude also systematically depended on stimulus size. Specifically, the remaining curves of Fig. 2A, B, D, E, G, H show eye position traces from three additional stimulus sizes that we used in our experiments, covering stimulus radii larger than approximately 1 deg. In all cases, the drift response was rendered larger with larger stimuli. When we now considered all of our tested stimulus sizes, we found that in both monkeys A and F, stimulus sizes beyond a radius of about 1-2 deg systematically, and monotonically, increased the amplitude of the drift response. In monkey M, this monotonic relationship was evident even from the very smallest stimulus sizes that we tested, well below 1 deg in radius. This latter observation can be better appreciated from Fig. 2C, F, I, summarizing the relationship between drift response magnitude and stimulus size. In these panels, and for each animal, we measured the drift response magnitude like we did in our earlier study (Malevich et al., 2020). Specifically, we took the difference in vertical eye velocity between two measurement intervals, a stimulus response epoch and a pre-stimulus baseline epoch (gray shaded regions in Fig. 1C; Materials and Methods). As can be seen from Fig. 2C, F, I, there was clear size tuning of the drift response magnitude in each animal: monkeys A and F showed a plateau (and even decreasing relationship in monkey A) up to about 1-2 deg, followed by a rise for larger stimuli; monkey M (generally having significantly faster baseline drift speeds) exhibited a monotonic increase with stimulus size, even for stimuli smaller than 1 deg in radius.

We confirmed the above interpretations statistically. We performed, within each animal’s data, a 1-way ANOVA relating drift response magnitude to stimulus size. In all three monkeys, there was a significant main effect of stimulus size [p<0.0001 for monkeys A, F, and M; F(7,6856) = 63.23, F(7,2331) = 57.78, and F(7,3205) = 50.71 for monkeys A, F, and M, respectively]. Therefore, besides still occurring for tiny foveal stimuli, the drift response also clearly exhibits size tuning, which we will later link to the size tuning of saccadic inhibition that we recently characterized in the same experiments (Khademi et al., 2023).

It is also interesting to note that in all three animals, larger stimulus sizes also increased the likelihood of observing a small transient modulation of eye position right at the very beginning of the overall drift response. For example, for the largest flashes, all three monkeys exhibited a small, but short-lived, downward change in eye position before the upward drift pulse (Fig. 2B, E, H, largest stimulus size), and this is similar to the downward transient that is evident in Fig. 1B. We frequently observed this small transient in our earlier study as well (Malevich et al., 2020). Monkey F additionally showed transient small oscillations in eye position at the beginning of the drift response for different sizes.

The larger stimuli in the current experiment additionally increased the likelihood that the upward drift response had a horizontal component to it. For example, monkey A’s upward drift response for large stimuli was accompanied by a slight leftward trajectory (Fig. 2A), and monkey M’s upward drift response for large stimuli was accompanied by a rightward trajectory (Fig. 2G). Once again, we observed such horizontal deviations accompanying the upward drift response in our earlier experiments as well (Malevich et al., 2020).

Therefore, our results so far demonstrate that the stimulus-driven ocular position drift response (Malevich et al., 2020) can still happen for tiny foveal visual transients, and that this drift response also exhibits size tuning (Fig. 2). As we will show below in more detail, it is interesting to note how this size tuning might relate to the size tuning of saccadic inhibition (Khademi et al., 2023).

### The drift response is stronger for high contrast stimuli

We next turned our attention to the contrast sensitivity curve of the drift response. We had the three monkeys view brief, transient full-screen flashes while they fixated their gaze at the center of the display. Across trials, the flashes (which were all darker than the background) could have a different Weber contrast (Materials and Methods). In all three animals, the drift response magnitude monotonically increased with stimulus contrast, increasing quasi-linearly as a function of log-contrast. These results can be seen in Fig. 3, which is organized similarly to Fig. 2. Specifically, Fig. 3A, B, D, E, G, H shows horizontal and vertical eye position traces from all three monkeys for three example contrast levels. The lowest tested contrast (5%; yellow curves) still showed a reliable drift response in all three monkeys. Moreover, the drift response magnitude increased with increasing contrast.

**Figure 3.**
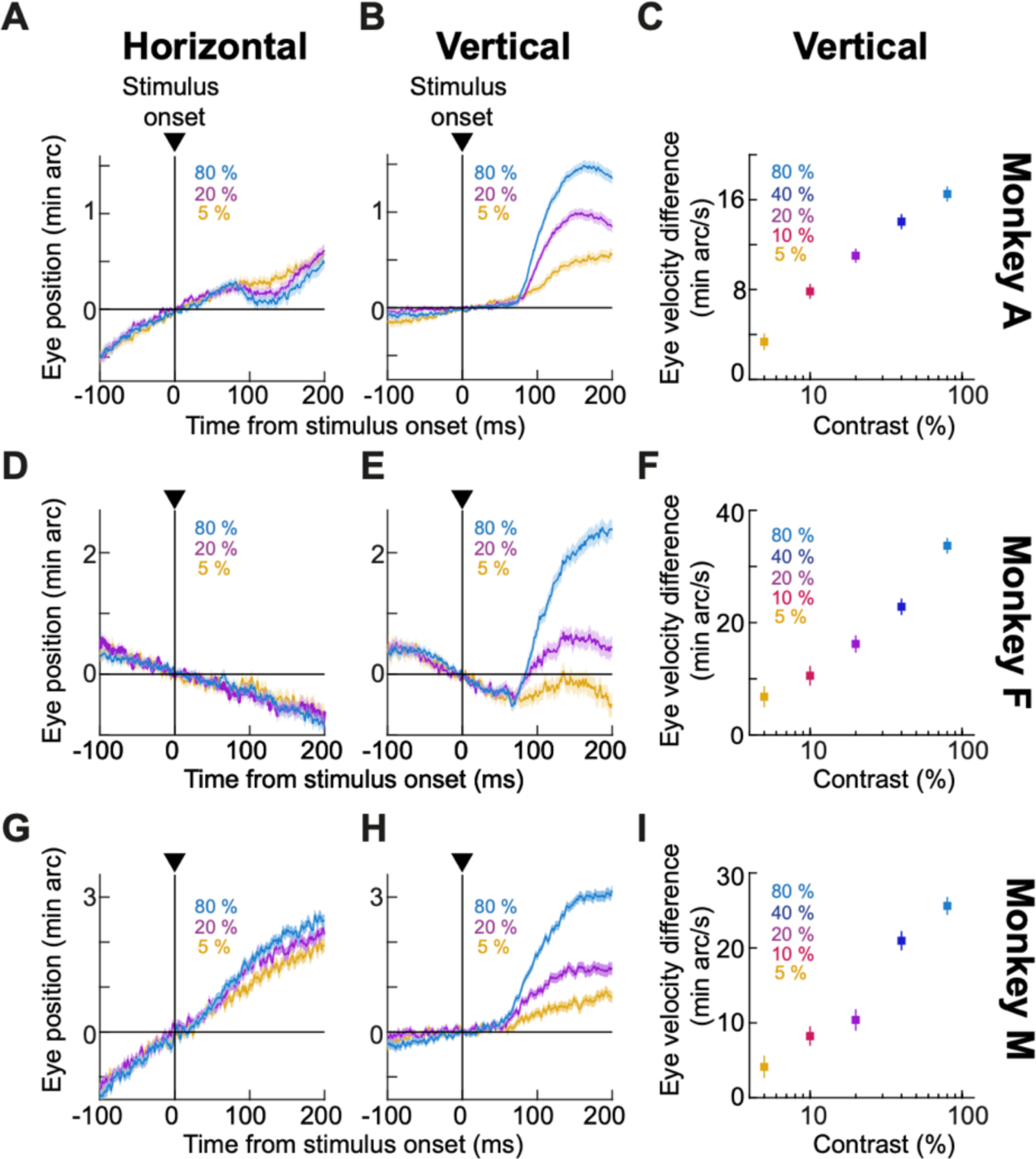
Contrast sensitivity of ocular position drift responses. **(A)** Average horizontal eye position from monkey A for three example stimulus contrasts (5%, 20%, and 80%). Error bars denote SEM, and the numbers of trials were 689, 739, and 750 for the three contrasts, respectively. **(B)** Average vertical eye position from the same trials as in **A** (error bars again denote SEM). A clear dependence of the ocular position drift response on contrast can be seen. **(C)** Our measure of the drift response magnitude for all tested stimulus contrasts in monkey A (n = 689, 699, 739, 754, and 750 trials from the lowest to the highest contrast). Error bars denote SEM. **(D-F)** Similar results for monkey F (n = 135, 165, 179, 223, and 262 trials from the lowest to the highest contrast). **(G-I)** Similar results for monkey M (n = 384, 412, 443, 433, and 475 trials from the lowest to the highest contrast). The figure is otherwise organized as Fig. 2.

To summarize these results, we again calculated the drift response size as described above (difference in vertical eye velocity between a response and a baseline epoch; Materials and Methods), and we plotted it as a function of stimulus contrast for each animal. These plots are shown in Fig. 3C, F, I, and they demonstrate the contrast sensitivity curve of the drift response. Statistically, there was a clear effect of contrast on drift response magnitude in each animal [p<0.0001 across all animals; 1-way ANOVA on drift response magnitude as a function of contrast; F(4,3626) = 56.65, F(4, 959) = 46.71, and F(4, 2142) = 45.43 for monkey A, F, and M, respectively].

Therefore, to the extent that stimulus-driven neural responses somewhere in the visual/oculomotor system might mediate short-latency ocular position drift responses (Malevich et al., 2020), these visual responses are expected to monotonically depend on stimulus contrast. Given the short time interval between stimulus onset and the actual eye movement modulations, we hypothesize (Buonocore and Hafed, 2023; Khademi et al., 2023) that these visual responses that are relevant for the drift response can be observed late in the oculomotor control circuitry, perhaps even in the brainstem pre-motor network.

#### The drift response is predominantly upward even for lower visual field stimuli

Speaking of oculomotor control circuitry, a candidate brain structure possessing short-latency visual responses and having direct access to the oculomotor system is the SC, and it is also a structure that can contribute to smooth eye movements (Krauzlis et al., 1997; Basso et al., 2000; Krauzlis et al., 2000; Hafed et al., 2008; Hafed and Krauzlis, 2008). Because the SC has stronger visual sensitivity for the upper visual field (Hafed and Chen, 2016; Fracasso et al., 2023), and seems to also magnify its representation for the upper visual field (Hafed and Chen, 2016), we hypothesized earlier that the predominantly upward nature of the drift response (for stimuli spanning both the upper and lower visual fields) might be mediated, at least partially, by SC visual activity (Malevich et al., 2020). If so, then presenting stimuli exclusively in the lower visual field (below the line of sight) should make the drift response downward instead, since it now shifts the balance of SC visual activity in favor of the lower visual field. We, therefore, next tested how the drift response was affected by presenting a half-screen brief flash either only in the upper half of the entire display or in the lower half (Materials and Methods). We also interleaved sham trials (without any flashes) as well as trials with small, localized flashes in the periphery (Materials and Methods). We note here that our earlier half-screen experiments (Malevich et al., 2020) involved splitting the screen area along the horizontal rather than vertical dimension (giving rise to either right or left visual field stimulation rather than upper/lower visual field stimulation); thus, these experiments still contained equal stimulus energy in the upper and lower visual fields and could not conclusively test the original hypothesis about upper visual field SC preference.

The drift response was still predominantly upward even for lower visual field half-screen stimuli. Figure 4 shows the eye position and velocity measures from this experiment in a manner similar to how we presented data in the earlier figures (Figs. 2, 3). The critical comparison here is between the upper and lower visual field stimulus conditions (red and purple colors in Fig. 4). In these conditions, the brief flash could consist of a black rectangle covering either exactly the top half or bottom half of the display. In each monkey, the drift response was still predominantly upward for lower visual field flashes (Fig. 4B, E, H), which is inconsistent with the hypothesis that SC visual responses dictate the upward direction of the drift response. Moreover, across the animals, there was no systematic relationship between the strength of the upward drift response and the visual field location of the stimulus. For example, in monkeys A and M, the overall drift response magnitude was similar for the upper and lower visual field stimuli (Fig. 4B for monkey A and Fig. 4H for monkey M). On the other hand, for monkey F, upper visual field stimuli did indeed cause a stronger upward component of the drift response than lower visual field stimuli (Fig. 4E). Statistical tests between the velocity difference measures of the two conditions confirmed these observations, as can be seen in Fig. 4C, F, I. In monkey A, there was no significant difference between upper and lower visual field flashes in Fig. 4C (p=0.26, t-test, t = -1241). For monkey F, the drift response magnitude was significantly stronger for the upper visual field stimuli (p=0.0079, t-test, t = 2.6642; Fig. 4F). And, for monkey M, there was again no reliable difference between the upper and lower visual field stimuli (p=0.77, t-test, t = -0.2818; Fig. 4I).

**Figure 4.**
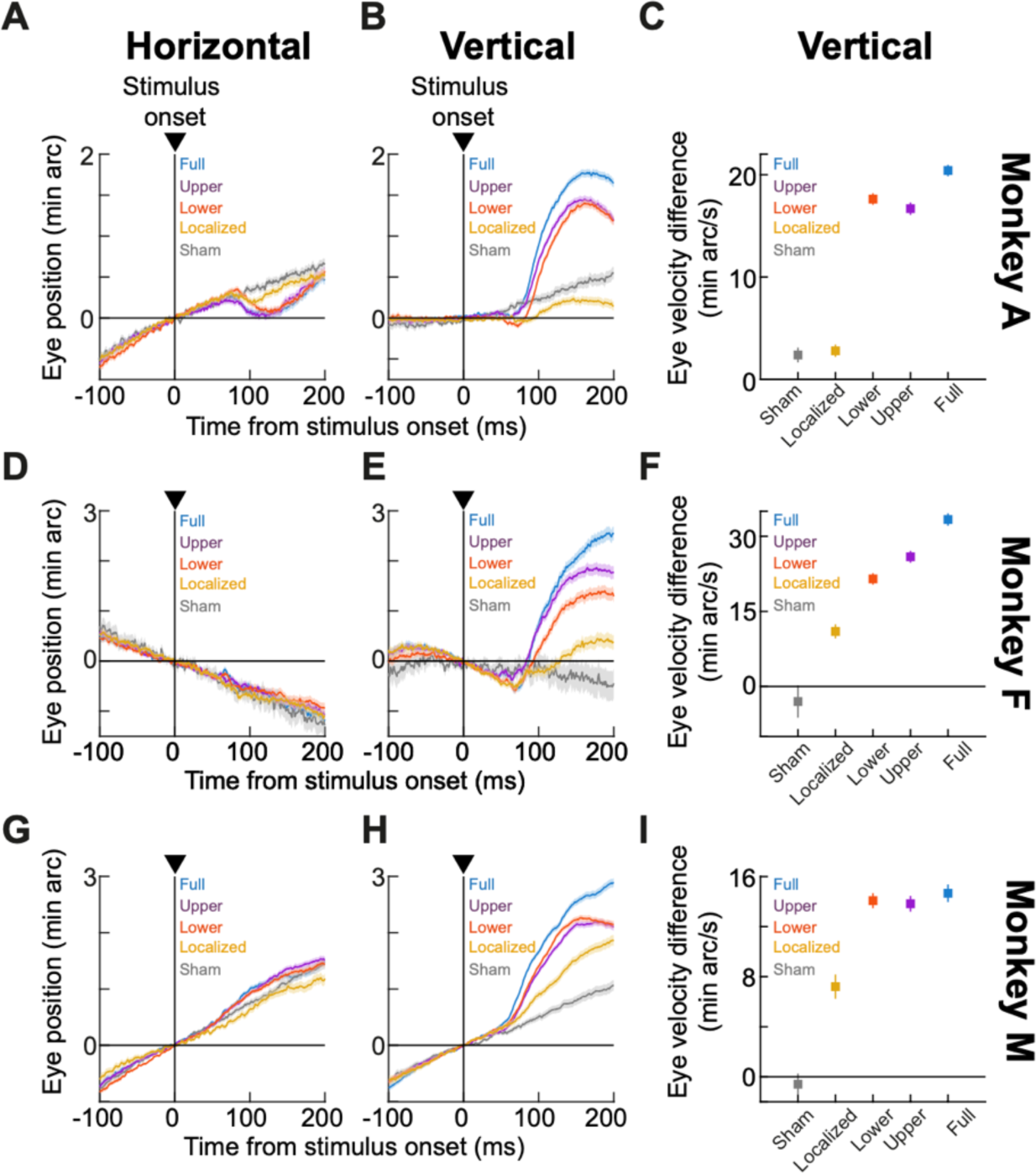
Predominantly upward ocular position drift responses even with lower visual field stimuli. **(A)** Average horizontal eye position from monkey A in the visual field experiment. Gray indicates sham stimulus onsets (n = 899 trials), yellow indicates a small localized flash eccentric from the fixation spot (Materials and Methods) (n = 833 trials), red indicates a stimulus onset in the lower half of the display (n = 890 trials), purple indicates a stimulus onset in the upper half of the display (n = 848 trials), and blue indicates a full-screen flash (n = 474 trials). Error bars denote SEM. **(B)** Average vertical eye position from the same trials (error bars again denote SEM). The drift response was predominantly upward even for lower visual field stimulus onsets (red). Note, however, how the initial downward component of the global drift response was weaker for the upper visual field stimulus onsets. **(C)** Our measure of the drift response magnitude for all conditions. Sham and localized stimulus onsets had weak drift responses (also see Figs. 5, 6); upper and lower visual field stimulus onsets had generally similar drift response magnitudes (and were both globally upward); and full-screen stimuli had stronger drift response magnitudes (consistent with the size tuning effects of Fig. 2). **(D-F)** Similar results for monkey F (n = 401, 341, 372, 415, and 72 trials for the shown conditions: sham, localized, lower visual field, upper visual field, and full-screen flashes, respectively). **(G-I)** Similar results for monkey M (n = 835, 439, 1208, 1143, and 553 trials). The figure is otherwise organized as Fig. 2.

Therefore, the drift response remains to be predominantly upward even with lower visual field stimuli, and the strength of this drift response may or may not reflect the presence of lower or upper visual field stimulus energy (also see later for further tests of this idea with small, localized flashes).

The other conditions shown in Fig. 4 were also informative in the broader context of this study. For example, in all animals, the drift response was always the strongest for the largest stimulus flashes (full-screen stimuli; blue colors in Fig. 4). This is consistent with our observations in Fig. 2. Interestingly, in the present experiments, we also interleaved trials with a 1 deg x 1 deg localized stimulus flash in the periphery relative to the fixation spot location (Materials and Methods; this is complementary to the small, foveal flashes of Fig. 2). Remarkably, there was still a small upward drift response in this case (all yellow curves in Fig. 4). This prompted us to investigate the influences of small, localized eccentric (rather than foveal) flashes on ocular position drifts in much more detail, as we describe next.

#### Small, localized stimuli additionally cause spatially-directed drift modulations

Our results so far demonstrate that the upward drift response occurs under a large variety of stimulus conditions, which hints that this drift response may be a reflexive movement of some kind. Indeed, the drift response remains predominantly upward even for lower visual field flashes (Fig. 4), and it also occurs for small foveal (Fig. 2) and eccentric (Fig. 4) targets. However, whether the drift response is a reflex or not, it is still likely the outcome of readout of stimulus-driven neural activity in the oculomotor control network. For small, localized targets, such activity can be highly spatially localized, especially in topographically organized structures like the SC. Might it then be the case that spatially localized visual bursts somewhere in the oculomotor system may play a modulatory role on ocular position drifts during fixation? Indeed, we recently found that at the time of saccade readout, spatially localized SC spiking systematically altered saccade metrics and kinematics even when such spiking was not part of the movements’ motor bursts (Buonocore et al., 2021), and the question now becomes whether a similar effect can be seen in ocular position drifts as well.

In previous work with peripheral cueing, we uncovered evidence that peripheral stimulus onsets can indeed give rise to spatially-directed drift trajectories (Tian et al., 2018), but our localized stimulus experiments in the drift response study of (Malevich et al., 2020) did not exhaustively study spatially-directed effects. Moreover, the stimulus locations for the localized targets in Fig. 4, and in (Tian et al., 2018), were not distributed enough to explore different spatially-directed modulations (Materials and Methods). Therefore, we explicitly ran an additional experiment with localized stimulus flashes, this time systematically sampling different directions relative to the line of sight.

The experiment consisted of the monkeys fixating a central spot, and a brief black flash of 1 deg x 1 deg size occurred at an eccentricity of 7.9 deg. The flash could occur at one of eight equally spaced directions relative to the fixation spot (see inset schematic in Fig. 5C). To robustly infer (from a statistical perspective) potential spatially-directed drift modulations, we first grouped all target locations along the horizontal direction. That is, any localized flash that was in the right visual field was grouped into the rightward target group, and any localized flash that was in the left visual field was grouped into the leftward target group (see the two different colors in the schematic inset of Fig. 5C). We then analyzed the eye positions of the three animals in the two groups of trials. We focused, here, on eye positions rather than eye velocities (like we did in earlier analyses) because we wanted to directly assess the potential spatial biasing that was caused by the stimulus onsets.

**Figure 5.**
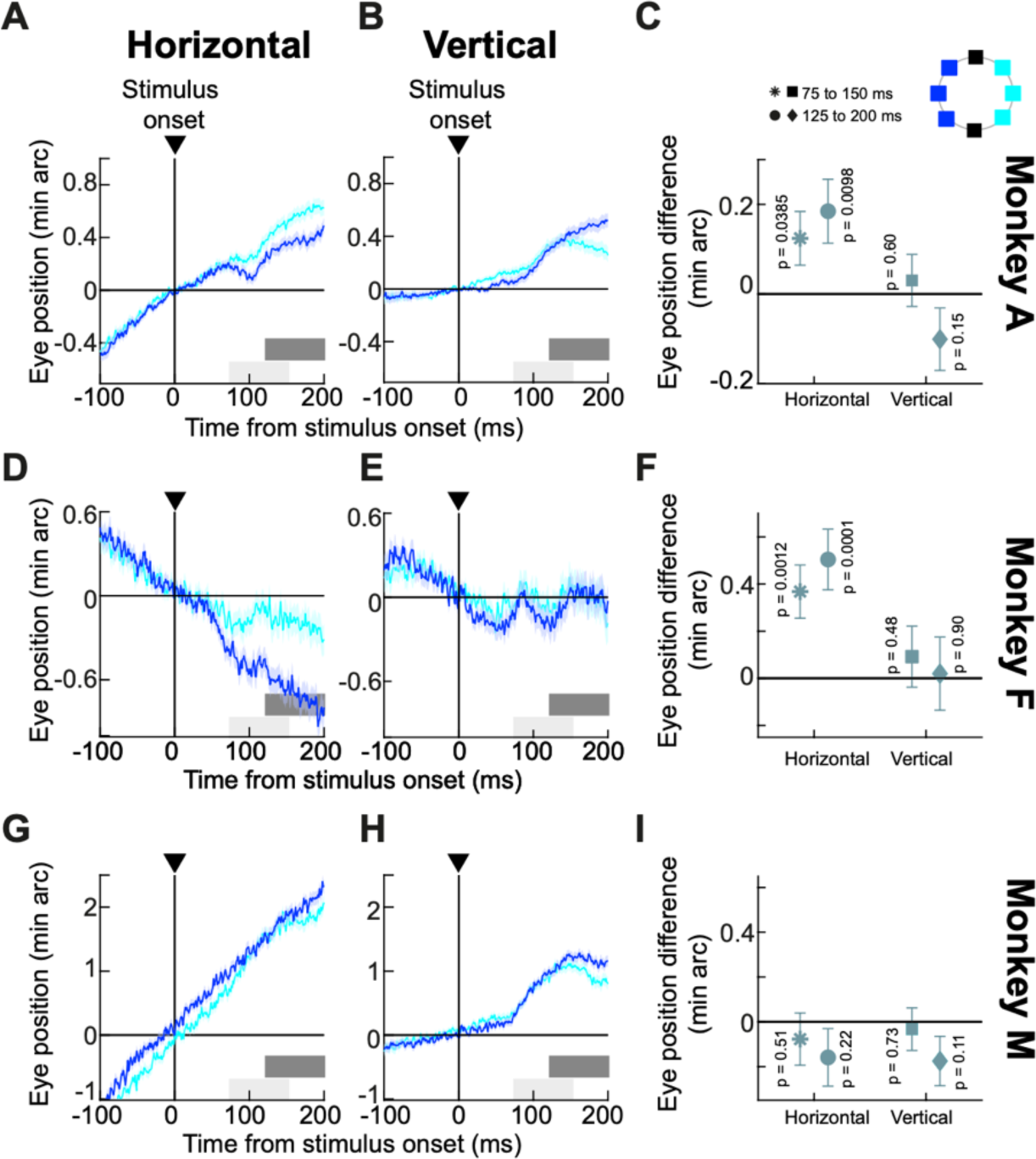
Spatially-directed drift modulations with localized stimuli along the horizontal direction. **(A)** Average horizontal eye position from monkey A when localized flashes (1 x 1 deg squares; 7.9 deg eccentricity) appeared in the right (cyan) or left (blue) visual field (see inset schematic in **C**). Error bars denote SEM (n = 1120 and 999 trials for right and left stimulus locations, respectively). Drift trajectory was affected by stimulus location, and the effect increased with time. The two gray bars near the x-axis indicate measurement intervals for comparing eye positions between the two groups of flash locations. **(B)** Vertical eye position from the same trials as in **A**. There was a general upward drift component, which was similar for rightward or leftward flashes. **(C)** We measured the difference between the cyan and blue curves in **A**, **B** for the two measurement intervals. Positive values mean rightward or upward differences between the cyan and blue curves. Horizontal eye position reflected the spatial layout of the flashes, and this difference increased with time. Vertical eye position did not. **(D-F)** Similar observations for monkey F (n = 349 and 398 trials for the right and left stimulus locations, respectively). This monkey showed an even clearer drift response modulation by stimulus location, also consistent with the same monkey’s performance in earlier experiments (Tian et al., 2018). **(G-I)** Similar analyses for monkey M (n = 649 and 1091 trials for the right and left stimulus locations, respectively). This monkey did not show horizontal modulation of drifts by stimulus location, but this monkey also had significantly faster baseline drift speed than the other two monkeys. As with the other two monkeys, there was still an upward stimulus-triggered drift response component (**H**). P-values indicate results of t-tests comparing eye positions within a given measurement interval.

Horizontal eye position drifts systematically reflected the peripheral hemifield locations of the brief, localized flashes, confirming our earlier observations that ocular position drifts can be spatially-directed (Tian et al., 2018). For example, Fig. 5A shows the horizontal eye position of monkey A for the two groups of stimulus locations (see inset schematic in Fig. 5C). As in all of our other analyses, we aligned eye positions at time zero to better appreciate the stimulus-driven changes in drift statistics. Shortly after stimulus onset, the monkey’s horizontal eye position deviated more rightward for the rightward flashes than for the leftward flashes, and the eye position deviation between the two stimulus groups increased in size with time. This modulation was riding on top of the upward drift response that we described above, as can also be seen from Fig. 5B. Here, the vertical eye position of the same animal and in the same trials showed an upward drift pulse, which (unlike horizontal eye position) was largely not differentiating between stimulus locations (especially in the early phases of the response). Thus, small, localized eccentric targets along the horizontal direction were associated with both an upward drift pulse as well as horizontal modulation of ocular position drifts reflecting the horizontal locations of the targets.

We summarized these observations by measuring the eye position difference between the two curves of Fig. 5A or Fig. 5B at two different post-stimulus times (shaded gray bars near the x-axes in Fig. 5A, B). This difference was significant for horizontal eye position but not for vertical eye position, as can be seen from Fig. 5C. Moreover, the horizontal difference in eye position was larger for the later time interval (Fig. 5C). These observations were virtually identical in monkey F (Fig. 5D-F), despite the monkey’s different baseline (pre-stimulus) drift trajectory. Thus, there can indeed be spatially-directed drift modulations in addition the upward drift pulse.

For monkey M, there was no clear evidence of spatially-directed drift modulations in the horizontal direction, but this monkey did exhibit a clear upward drift pulse (Fig. 5G-I). As mentioned earlier, this monkey had the fastest baseline drift speeds from among the three animals, rendering a weak modulation by spatially localized peripheral activity harder to see. This is similar to our observations of the size tuning experiments described above (Fig. 2).

In all, the results of Fig. 5 confirm that ocular position drifts are not always random or stochastic (Kowler and Steinman, 1979b, a; Ahissar et al., 2016; Tian et al., 2018; Skinner et al., 2019; Bowers et al., 2021; Reiniger et al., 2021; Clark et al., 2022; Nghiem et al., 2022), and that these drifts can reliably reflect localized stimulus locations in addition to exhibiting a (potentially reflexive) upward drift pulse. Having said that, true dependence of ocular position drifts on localized stimulus locations should include evidence of spatially-directed drift trajectories for the vertical dimension as well. Thus, we next regrouped our trials according to the vertical locations of the localized flashes (see inset schematic of Fig. 6C). In this case, all three monkeys showed evidence that vertical eye position deviated more upward for upper visual field target locations than for lower visual field target locations (Fig. 6); the effect was weakest in monkey A, but the trend was still clearly there. Moreover, in all cases except for monkey M, horizontal eye position deviations were similar to each other for the upper and lower visual field targets, exactly complementary to the results of Fig. 5. Thus, in Fig. 5, it was horizontal eye position that was most affected by horizontal target locations, and in Fig. 6, it was vertical eye position instead that was most affected by vertical target locations. Such a complementary nature of the results of Figs. 5, 6 is consistent with the interpretation that spatially-directed drift responses can indeed occur. Once again, these spatially-directed effects were occurring in addition to a global upward drift response, which was similar to what we saw in all of our earlier analyses with other types of stimuli.

**Figure 6.**
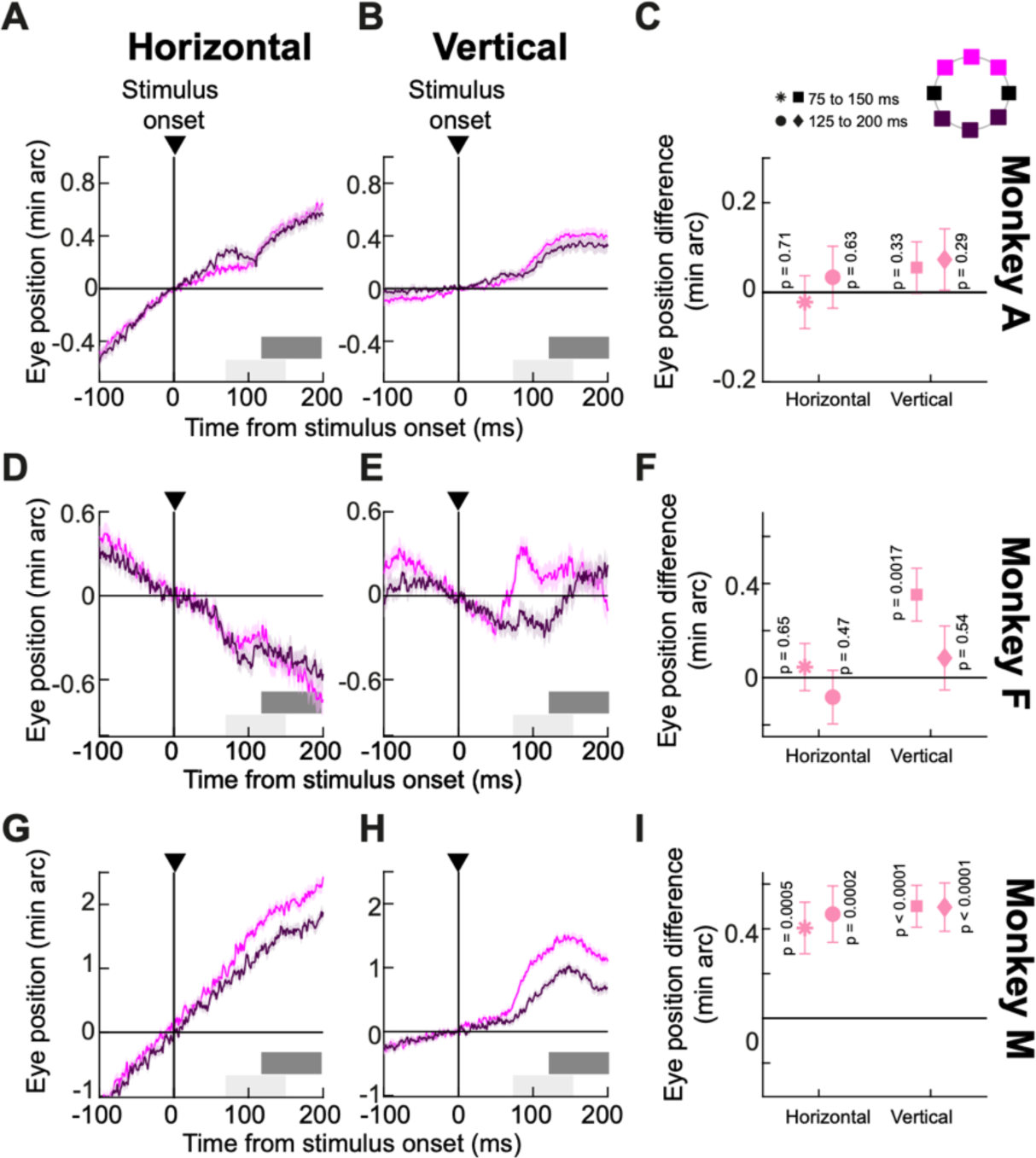
Spatially-directed drift modulations with localized stimuli along the vertical direction. This figure is organized exactly like Fig. 5, except that we now grouped the trials according to whether the localized stimulus flashes were in the upper or lower visual field (see inset schematic in **C** for the color codes). All monkeys showed a vertical drift response that was predominantly upward. On top of that, the stimulus locations now modulated the vertical component of eye positions more than the horizontal component, again consistent with the idea that localized stimuli can still have a modulatory effect on ocular position drifts (compare the eye position traces to those in Fig. 5). Also note that the vertical position difference measurements in the later time interval did not increase relative to those in the earlier time interval as in Fig. 5 for the case of horizontal position difference (in monkeys A and F). This is likely because the spatially-driven modulation in the vertical dimension was riding on a drift response that was already predominantly vertical in the current case. **(A-C)** n = 1100 and 1006 trials for upper and lower visual field stimulus locations, respectively. **(D-F)** n = 303 and 312 trials for upper and lower visual field stimulus locations, respectively. **(G-I)** n = 984 and 881 trials for upper and lower visual field stimulus locations, respectively.

Therefore, ocular position drifts exhibit a stimulus-driven upward drift response for a large range of stimulus types (including small foveal and peripheral targets; Figs. 1-4), and they also undergo spatially-directed modulations by spatially localized flashes (Figs. 5, 6). These spatially-directed modulations likely reflect localized visual bursts in oculomotor control circuits, such as the SC, that have an impact on eye movement generation in the brain. It would be interesting in the future to understand why large (non-spatially-specific flashes) in the upper and lower visual field (Fig. 4) did not systematically modulate the drift response in the vertical eye position direction across all three animals even though small targets did (compare the vertical eye position results of Fig. 4 and Fig. 6).

### The drift response is synchronized with saccadic inhibition

Our analyses so far focused on trials in which there were no saccades in the interval from - 100 ms to 200 ms relative to stimulus onset. This was important to allow us to best observe the drift response, because saccades would cause much larger velocity pulses that would mask such a response (but see our later analyses in which we directly tackled the question of saccade-drift interactions). We also know from our recent work (Malevich et al., 2020) that the drift response is complementary to saccade generation, in the sense that it occurs near the time of saccadic inhibition. Having said that, our current study afforded us a much better chance at exploring this complementary nature between saccade generation and the drift response in more detail. Specifically, we know from our most recent work that the time of saccadic inhibition in our size tuning and contrast sensitivity experiments varied systematically as a function of stimulus type (Khademi et al., 2023). If the drift response is indeed obligatorily synchronous to saccadic inhibition, then we should also see evidence that the timing of the drift response (not just its magnitude like in our earlier analyses above) should depend on the stimulus feature. This would, in turn, imply that the drift response and saccadic inhibition may be generated by common neural circuitry.

We explored this idea by plotting drift responses and saccades together in the same graphs, and we checked whether drift response timing co-varied with saccadic inhibition timing. Figure 7 illustrates this for the size tuning experiment. For each monkey, the individual rasters indicate individual saccade times across trials, grouped by stimulus size (different colors). These rasters were reproduced from our earlier study (Khademi et al., 2023), since we analyzed drift responses from the same set of experiments. Superimposed on the rasters, we additionally plotted average vertical eye positions for each stimulus size (similar to the example vertical eye position plots in Fig. 2). Each eye position curve was scaled to fit within the similar-colored group of saccade rasters, and position scale bars for each curve are included (on the left side of the curve) for reference. As can be seen, the drift response latency appeared synchronized with the latency of saccadic inhibition, as estimated by the L_50_ parameter (dark green vertical lines; Materials and Methods). This parameter is routinely used to characterize the latency of saccadic inhibition (Reingold and Stampe, 2002, 2004; Rolfs et al., 2008; Khademi et al., 2023), and Fig. 7 shows that when L_50_ was late, so was the onset of the drift response, and vice versa.

**Figure 7.**
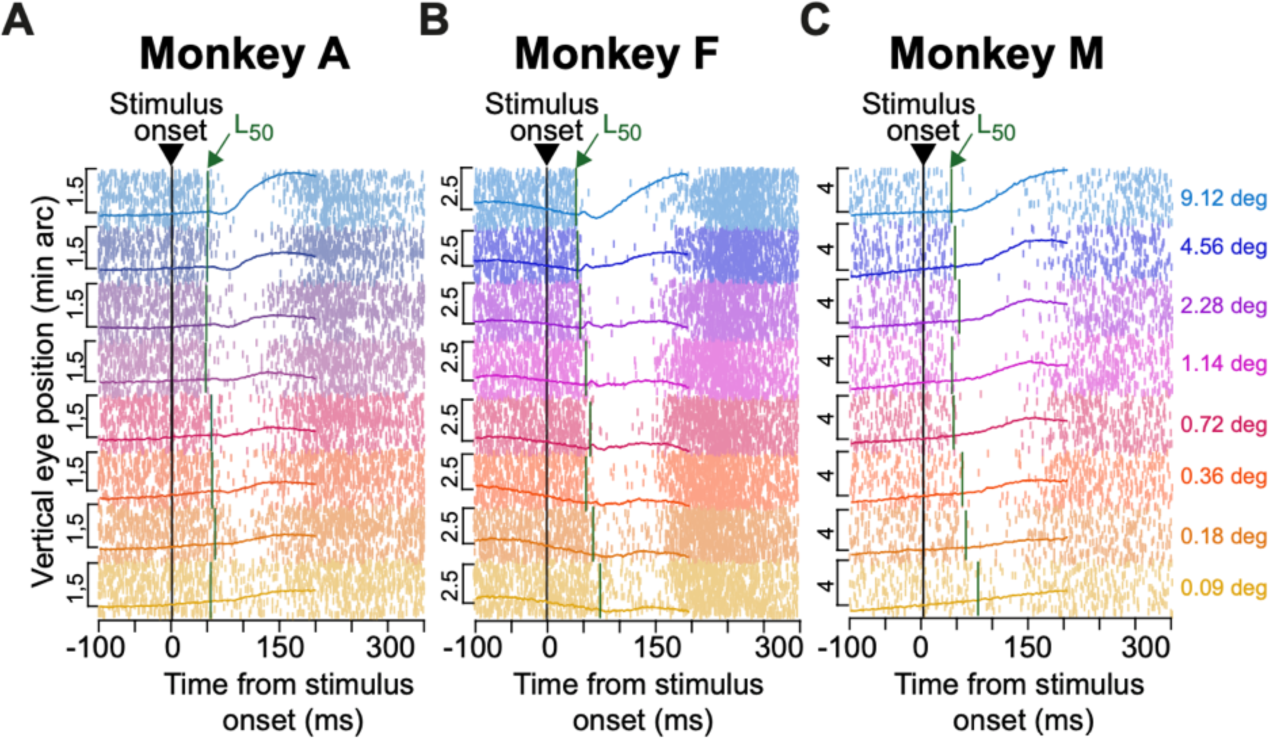
Coincidence between drift response onset and saccadic inhibition timing. **(A)** In our size tuning experiment, we recently found that the timing of saccadic inhibition depends on stimulus size (Khademi et al., 2023). This is indicated here, for monkey A, by the raw saccade onset times (tick marks) and a measure (vertical dark green lines marked with L50) of saccadic inhibition timing (Materials and Methods) (Khademi et al., 2023). Each row of tick marks represents a single trial, and each tick mark represents the onset time of a saccade. The L50 line in each condition (dark green color) indicates our estimate of the saccadic inhibition timing (Khademi et al., 2023), and all trials of a given stimulus size are grouped together according to the color legend. Within each group of trials, we also plotted the drift response (on trials without saccades; Materials and Methods) by showing vertical eye position aligned on stimulus onset (scale bars are shown on the left of each curve). Despite the variable saccadic inhibition timing, the drift response was synchronized with such timing. That is, both the timing of the drift response (on trials without saccades) and the timing of saccadic inhibition (on trials with saccades) depended on the stimulus properties (also see Figs. 8, 9). **(B)** Similar observations from monkey F. **(C)** Similar observations from monkey M. The saccade data in **B** were directly replotted from (Khademi et al., 2023) (CC-BY) since they came from the same experiments. Numbers of trials in the saccade data can be inferred from the rasters and from (Khademi et al., 2023); numbers of trials in the smooth drift data were reported in Fig. 2.

We next checked this synchrony idea further by asking whether our drift response curves across stimulus sizes were better aligned to stimulus onset or to the onset of saccadic inhibition. For each animal, we plotted in Fig. 8 the average vertical eye position traces for all stimulus sizes (the curves were displaced vertically from each other for easier viewing). In the top row of the figure (Fig. 8A, C, E), the traces were aligned to stimulus onset like in our earlier analyses, and the small vertical tick marks indicate the time of saccadic inhibition (L_50_) as we recently calculated it (Khademi et al., 2023). In the bottom row (Fig. 8B, D, F), the same traces were now aligned to the time of L_50_, with the small vertical tick marks now indicating stimulus onset time. In all three monkeys, the drift response curves were better synchronized with L_50_ than with stimulus onset. That is, the curves across the different stimulus sizes were less jittered in time relative to each other when they were referenced to L_50_ than to stimulus onset time. Thus, there seems to be an obligatory timing relationship between saccadic inhibition and drift response latency.

**Figure 8.**
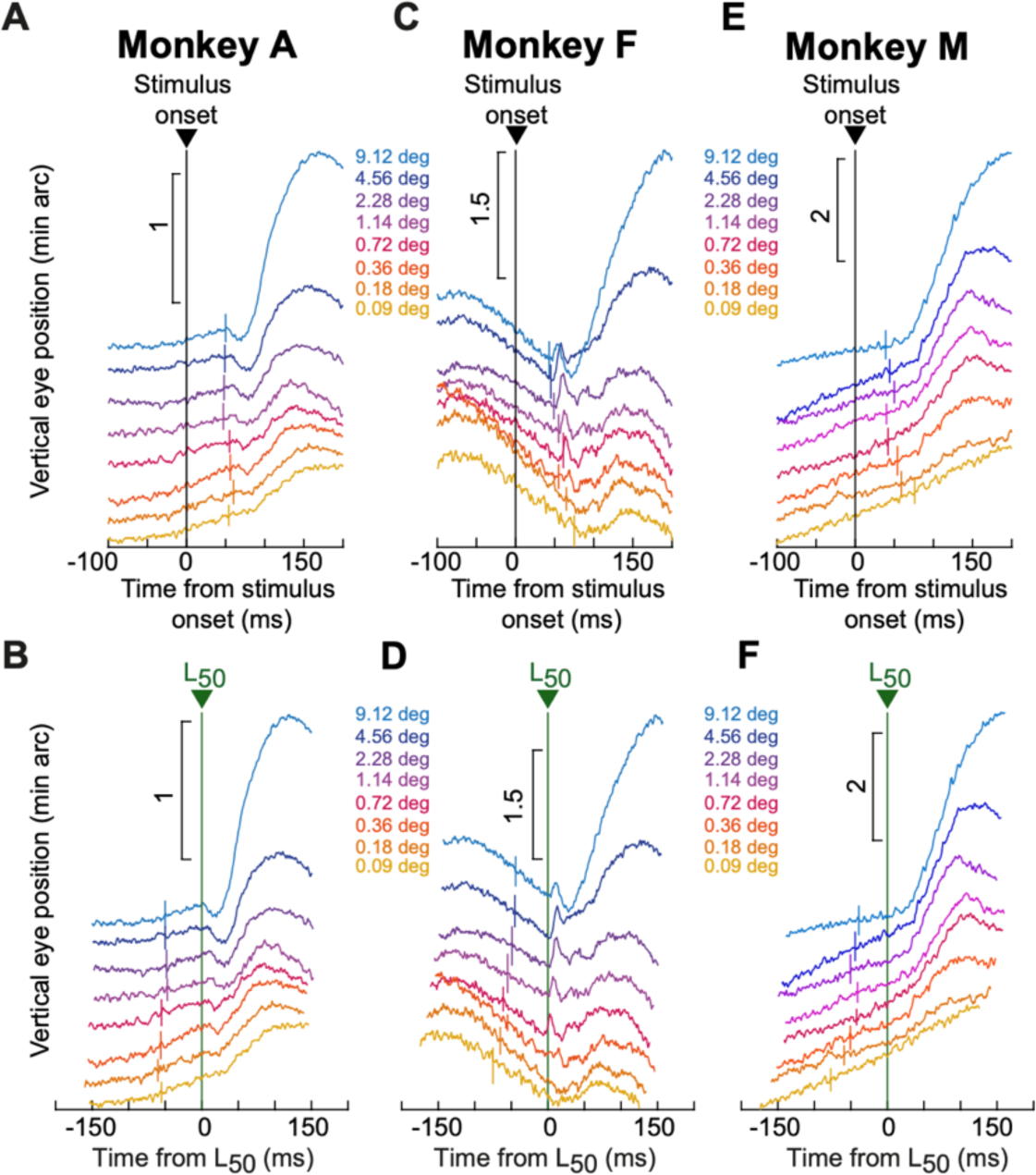
Alignment of the drift response onset to saccadic inhibition timing. **(A)** Average vertical eye position in each condition of the size tuning experiment from monkey A. Each curve was slightly offset vertically from the others for easier viewing. The vertical tick mark in each curve indicates the time of saccadic inhibition for the condition, as estimated by the parameter L50 (Materials and Methods) (Khademi et al., 2023). Consistent with Fig. 7, saccadic inhibition time varied with stimulus size (Khademi et al., 2023), and the drift response followed this relationship. **(B)** This is better seen when aligning the drift response curves of **A** to the time of L50 rather than to the time of stimulus onset. Here, all the curves were better aligned in time. The vertical tick marks now indicate stimulus onset time. **(C, D)** Similar results for monkey F. **(E, F)** Similar results for monkey M. In all cases, the drift response was relatively well synchronized with the timing of saccadic inhibition, potentially suggesting a common mechanism underlying both phenomena. The numbers of trials underlying each curve were reported in Fig. 2.

Such an obligatory relationship also held in our contrast sensitivity experiment. In this experiment, lower contrasts were generally associated with later saccadic inhibition (Khademi et al., 2023). As Fig. 9 shows, such contrasts were also associated with later drift responses, and across stimulus contrasts, the timing of the drift responses appeared to be better temporally aligned to the timing of saccadic inhibition across stimulus features (Fig. 9B, D, F).

**Figure 9.**
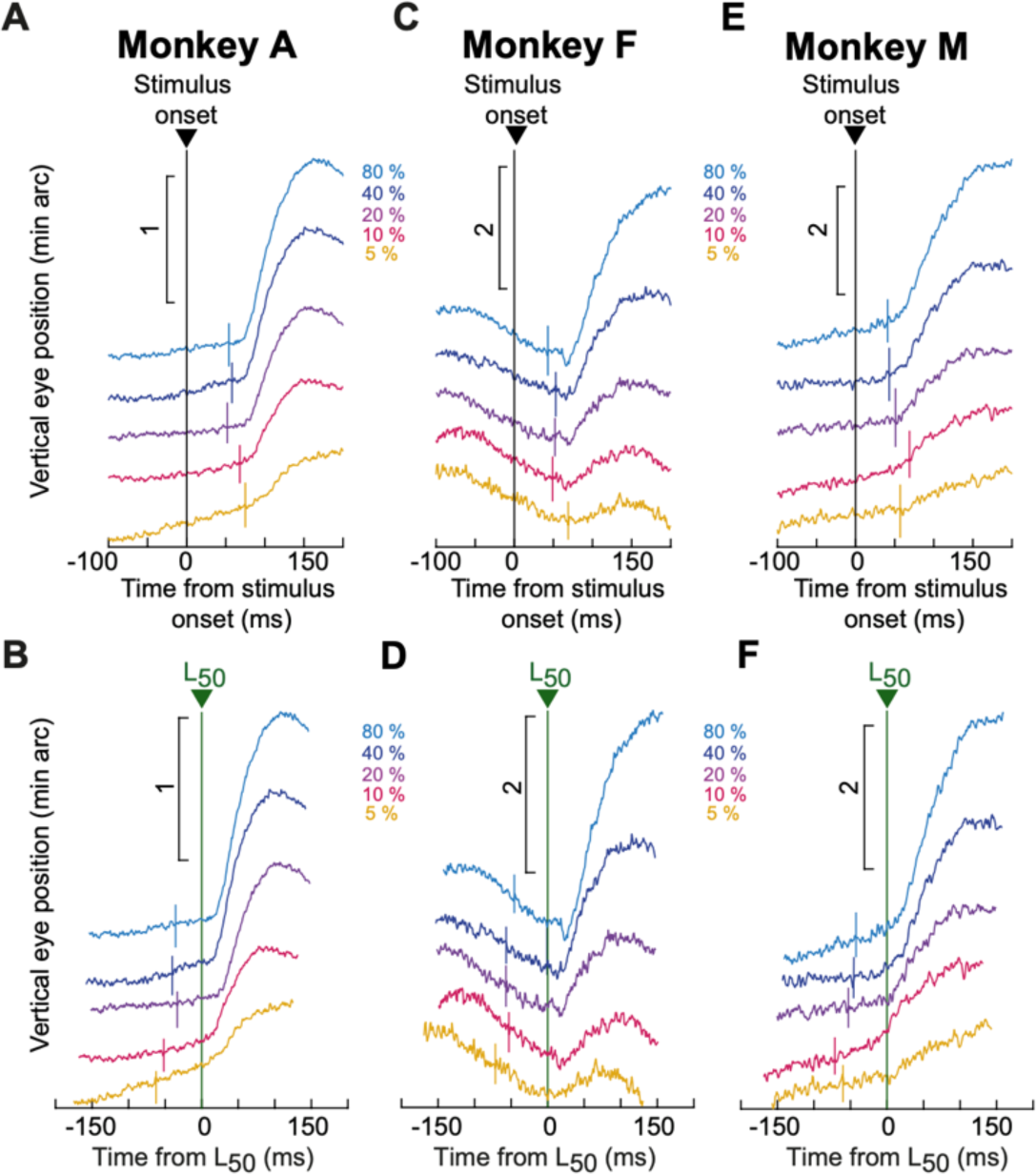
Alignment of the drift response onset to saccadic inhibition timing in another task. This figure is organized similarly to Fig. 8, but now showing results from the contrast sensitivity experiment. Once again, saccadic inhibition time depended on stimulus property (Khademi et al., 2023), and once again, the drift response was synchronized with the timing of saccadic inhibition. The figure is otherwise formatted identically to Fig. 8, and the numbers of trials underlying each curve were reported in Fig. 3.

Therefore, across multiple tasks associated with multiple different times of saccadic inhibition (Khademi et al., 2023), we found that the drift response was synchronized with the reflexive interruption of saccade generation rhythms caused by visual onsets in the environment.

### The drift response occurs with different starting eye positions

In addition to initially mentioning the potential relationship between the drift response and saccadic inhibition, we also suggested in our earlier work that the drift response occurs independently of starting eye position (Malevich et al., 2020). However, in that study, we only used the natural variability of eye positions during fixation to test whether the drift response still occurred when the eye was momentarily fixating below or above some central value (such as the median eye position across trials). This left open the question of whether the drift response might depend on significantly larger eye position deviations from the primary position. To answer this, we performed a new version of our contrast sensitivity experiment, in which we now explicitly required gaze fixation away from the display center. Specifically, in each block of trials, we placed the fixation spot at 4 deg eccentricity from the center of the display, either to the right of it, to the left of it, above it, or below it (Fig. 10A).

**Figure 10.**
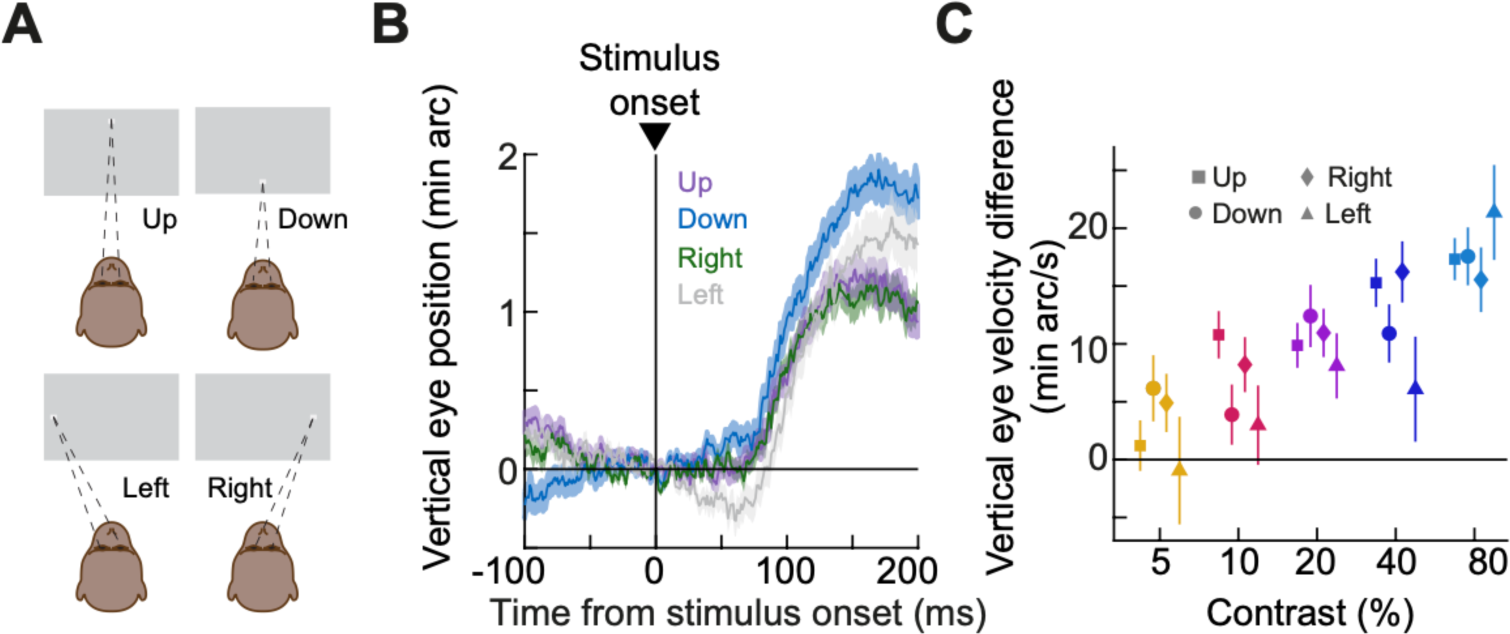
Independence of the drift response from starting eye position. **(A)** We performed the contrast sensitivity experiment, but now requiring gaze fixation at 4 deg eccentricity from the center of the display (either to the right, left, up, or down from display center). **(B)** Average vertical eye position from the four conditions with the highest contrast stimulus (error bars denote SEM, and n = 62, 71, 58, and 55 trials for the up, down, right, and left gaze fixation conditions, respectively). The upward drift response always occurred, even when the eye was gazing down. Note that the pre-stimulus drift direction showed some dependence on gaze position. For example, downward gaze position was associated with more upward pre-stimulus eye position drift, whereas upward gaze position was associated with more downward pre-stimulus eye position drift (compare the blue and purple curves). However, in both cases, the stimulus-driven response was still upward. **(C)** Our measure of the drift response magnitude as a function of stimulus contrast and fixation gaze position. The drift response was stronger with higher contrasts. However, there was no systematic dependence on gaze position – a two-way ANOVA revealed a significant main effect of stimulus contrast [F(4,1184) = 16.42; p<0.0001] but not starting eye position [F(3,1184) = 1.36; p = 0.25]. This extends our earlier findings with much smaller starting gaze position deviations (Malevich et al., 2020). The numbers of trials per condition were as follows: 63, 66, 44, and 49 for up, down, left, and right, respectively (5% contrast); 60, 73, 45, and 59 for up, down, left, and right, respectively (10% contrast); 61, 71, 52, and 60 for up, down, left, and right, respectively (20% contrast); 61, 78, 55, and 49 for up, down, left, and right, respectively (40% contrast); 62, 71, 58, and 55 for up, down, left, and right, respectively (80% contrast).

In all cases, the drift response still occurred, and it was largely independent of the starting eye position. Figure 10B shows vertical eye position traces for the highest contrast stimulus from each gaze position condition. Of course, and as with all of our earlier analyses, we aligned all traces to the eye position at stimulus onset, and that is why all curves are aligned to zero eye position on the y-axis despite the different starting gaze position conditions. As can be seen, the upward drift response always happened, irrespective of starting eye position. Interestingly, the pre-stimulus drift trajectory did depend on gaze position. For example, when gaze was up (purple curve), pre-stimulus drift in vertical eye position was downward, and when gaze was down (blue curve), pre-stimulus drift in vertical eye position was upward. Nonetheless, and as just stated, there was still an upward drift response in both cases.

Across all stimulus contrasts, we replicated the contrast sensitivity curve of Fig. 3 for each gaze position condition (Fig. 10C). Indeed, there was no effect of gaze position on drift response magnitude, but there was a clear effect of stimulus contrast; statistical results are presented in the legend of Fig. 10. Therefore, even with substantial deviations of gaze positions, the drift response still occurs, and it is still predominantly upward. Moreover, pre-stimulus drift trajectories can depend on gaze position, likely reflecting a pulling force (whether biomechanical or neural) to return the eye back to the primary position. Nonetheless, relative to these changed baseline drift statistics, the drift response magnitudes are more-or-less constant (Fig. 10C).

### The drift response magnitude is affected by the occurrence of peri-stimulus saccades

Finally, and still on the general theme of interactions with saccades (Figs. 7-9) and gaze positions (Fig. 10), we next explored modulations in the drift response magnitude by the occurrence peri-stimulus saccades. In our earlier work (Malevich et al., 2020), a coarse analysis suggested minimal (or even potentially no) interaction with peri-stimulus saccades. However, due to data sparsity, the analysis that we conducted at the time was not specific enough in its time course resolution. For example, rather than testing trials with saccade onsets occurring within only a constrained time interval (as we would typically do for studying transient modulations by saccades), we tested trials with “saccades up to” some particular time point. Such an analysis might have excessively blurred transient changes in drift response magnitude caused by the occurrence of peri-stimulus saccades (indeed, peri-saccadic effects can be very transient in nature). With our current experiments, we had an opportunity to explore such transient changes in more detail. Indeed, because suppression of both visual sensitivity and perception by peri-stimulus saccades is jumpstarted already in the retina (Idrees et al., 2020; Idrees et al., 2022), it would be remarkable if the drift response magnitude was completely unaffected by saccades. This would suggest that whatever visual response is mediating the drift response would be immune to peri-saccadic suppression. This question, therefore, warranted more detailed analysis in the current study.

Here, we binned our data for investigations of potential “saccadic suppression” as we usually do for analyzing visual neural sensitivity (Hafed and Krauzlis, 2010; Chen and Hafed, 2017; Fracasso et al., 2023) or perception (Idrees et al., 2020; Baumann et al., 2021). For example, for a given stimulus condition, we took all trials in which there was a saccade onset occurring within the interval between -100 ms and 0 ms relative to stimulus onset (green shaded region in Fig. 11A). These trials would be expected to exhibit suppressed visual sensitivity if saccadic suppression does take place. We also took trials in which there was a saccade onset 175-275 ms after stimulus onset (yellow shaded region in Fig. 11A). These trials, instead, would be expected to not experience saccadic suppression (since the saccades occurred far away in time from stimulus onset). Finally, we took trials in which there were no saccades at all in the interval from -100 ms to 200 ms relative to stimulus onset (shaded gray region in Fig. 11A), and these trials constituted our “standard” drift response trials (like in our other analyses above).

**Figure 11.**
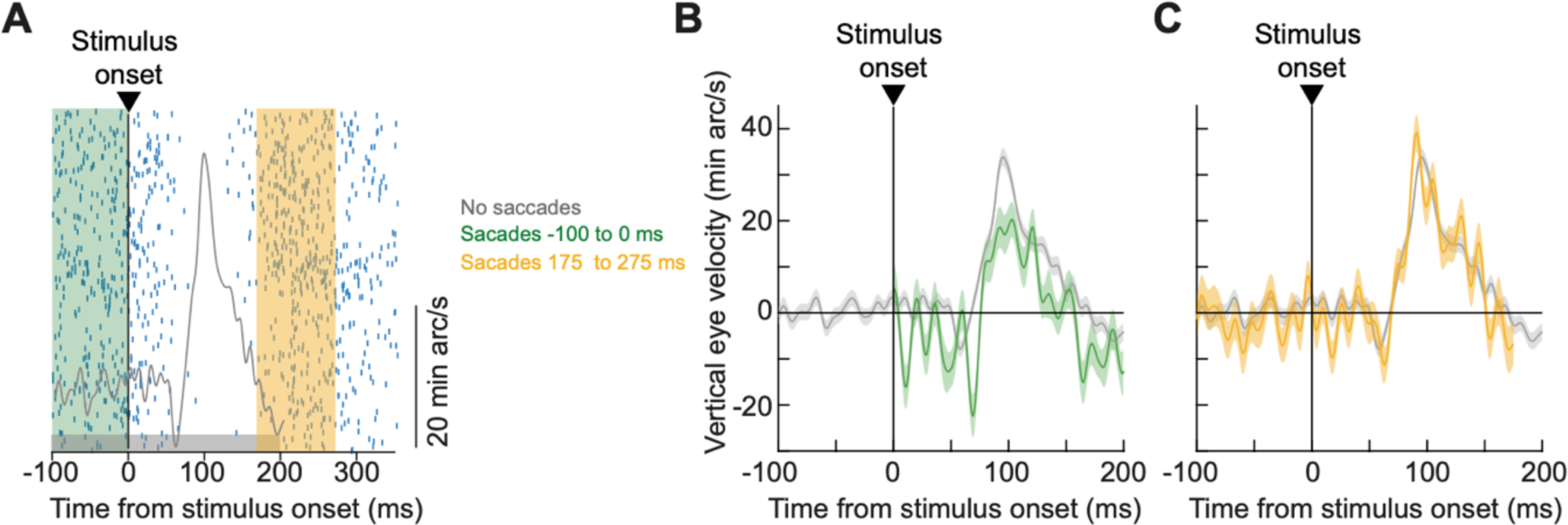
Saccadic suppression of drift responses. **(A)** Example saccade raster plot and drift response (shown by vertical eye velocity) from one monkey (A) and one condition (9.12 deg radius in the size tuning experiment). The shaded colored bars indicate how we picked trials to check for an interaction between peri-stimulus saccades and drift responses. For each such bar, we picked only trials from the same condition having saccade onsets occurring within the bar’s time window. The shaded gray bar, on the other hand, indicates our standard approach to analyze no-saccade drift responses. Note that we did not sample all peri-stimulus saccade times with high resolution; this was done to increase robustness of our observations, especially given how noisy velocity measures can be with small numbers of trials. Nonetheless, we had sufficient data to check whether stimulus onsets immediately after nearby saccades (shaded green interval) had altered drift responses. **(B)** For such trials (green), the drift response magnitude was suppressed. Error bars denote SEM (n = 168 and 879 for the green and gray curves, respectively), and no eye velocity data are shown in the green curve in the interval from -100 to 0 ms because saccades were occurring. As with the case of saccadic suppression (Hafed and Krauzlis, 2010; Chen and Hafed, 2017), the drift response was suppressed, suggesting that it might depend on circuits in which visual responses experience saccadic suppression; note that this observation was also categorically different from post-saccadic enhancement (Chen and Hafed, 2013). **(C)** For trials with a saccade occurring 175 to 275 ms after stimulus onset (well away from stimulus onset), the drift response was recovered. Error bars again denote SEM (n = 171 and 879 for the colored and gray curves, respectively). Also see Fig. 12 for summary data of suppression and recovery across other conditions and tasks.

The drift response magnitude was suppressed by the presence in peri-stimulus saccades. In Fig. 11B, for an example monkey and condition, we compared the standard drift response (gray curve in both panels A and B of Fig. 11) to the response when the stimulus occurred right after microsaccades during pre-stimulus fixation (green). As can be seen, the upward stimulus-evoked velocity pulse was smaller in peak amplitude when the microsaccades occurred than when they did not occur. On the other hand, for microsaccades distant in time from stimulus onset (yellow in Fig. 11), the drift response was recovered (Fig. 11C). Thus, for a brief moment in time when stimulus onset occurred near saccade onset, the subsequent stimulus-driven drift response was systematically suppressed. This is qualitatively very similar to the classic phenomenon of saccadic suppression.

This observation was consistent across all monkeys and in all conditions that we checked. For example, for each stimulus condition in both the contrast sensitivity (5 stimulus conditions) and size tuning (8 stimulus conditions) tasks, we measured the drift response magnitude (as we did earlier; Figs. 2-4, 10) and plotted it as a function of which time window of Fig. 11A the particular trials came from. For trials with saccades -100-0 ms from stimulus onset, the drift response magnitude was always smaller than the drift response magnitude in the absence of peri-stimulus saccades (Fig. 12; compare the response in the peri-stimulus time bin centered on -50 ms to the corresponding baseline response and its associated horizontal dashed line). Moreover, for trials with saccades 175-275 ms from stimulus onset, the drift response magnitude was recovered and much closer to the standard drift response magnitude in the absence of peri-stimulus saccades (Fig. 12; compare the response in the later time bin to that in the associated horizontal dashed line). We also confirmed these observations statistically. For example, a two-way ANOVA in the contrast sensitivity task revealed a main effect of both stimulus contrast [p<0.0001 in monkeys A, F, and M] and saccade time relative to stimulus onset [p<0.0001 in monkeys A, F, and M]. There was also a significant interaction between saccade time and stimulus contrast in monkey A [F(4,1343) = 3.76; p = 0.0048] but not in either monkey F [F(4,1744) = 0.54; p = 0.70] or monkey M [F(4, 1162) = 0.89; p = 0.47]. Similarly, a two-way ANOVA in the size tuning task revealed a main effect of both stimulus radius [p<0.0001 in monkeys A, F, and M] and saccade time [p<0.0001 in monkeys A, F, and M] in all three monkeys. However, once again there were no consistent interaction effects. Monkey A showed no significant interaction between stimulus radius and saccade time [F(7,2633) = 1.38; p =0.21], monkey F showed a significant interaction [F(7, 4118)= 5.17; p < 0.0001], and monkey M showed no significant interaction [F(7,1542) = 1.7; p = 0.11].

**Figure 12.**
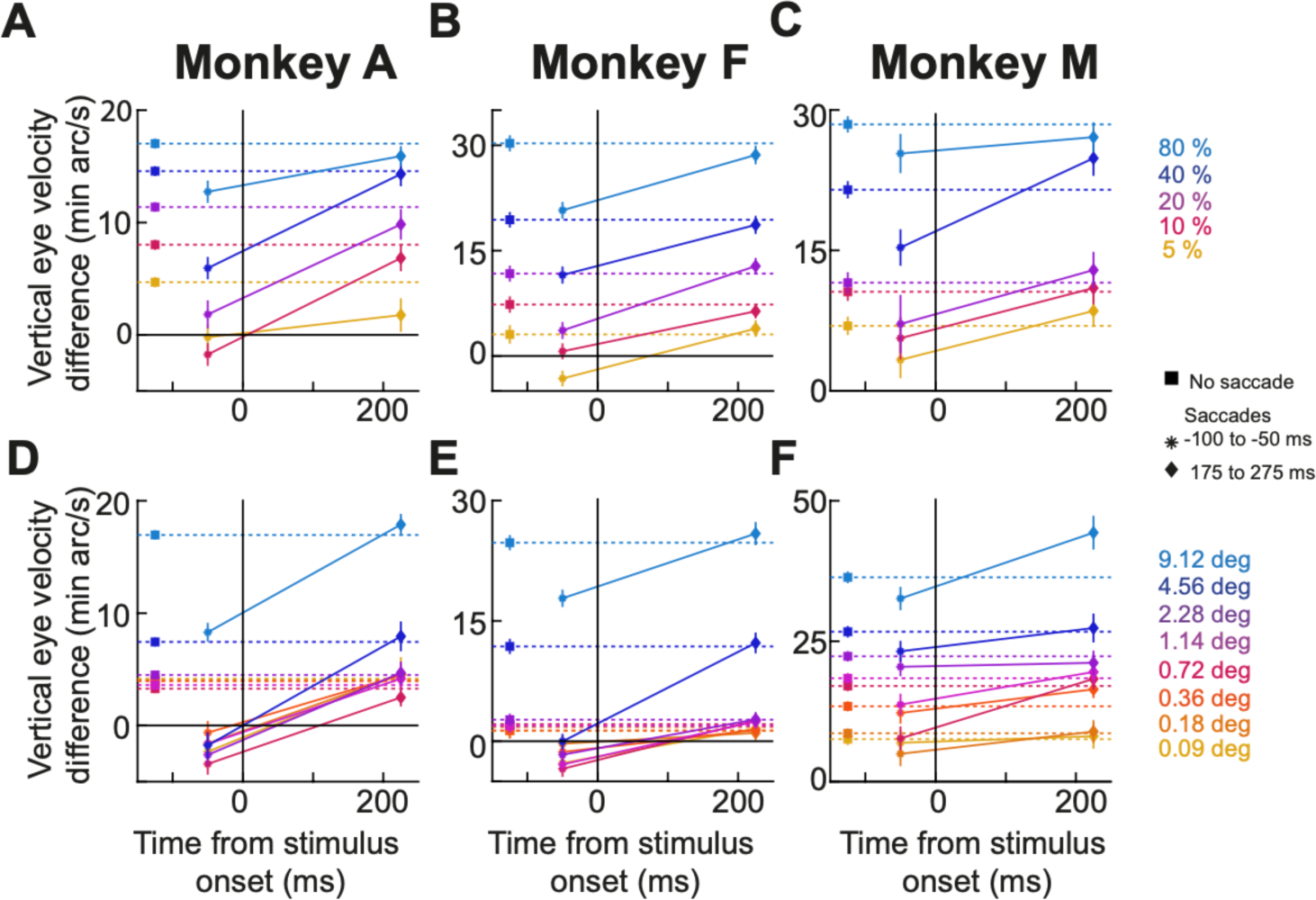
Suppression of the drift response strength by the occurrence of peri-stimulus saccades. **(A-C)** Summary plots of saccadic suppression of the drift response strength for each monkey in the contrast sensitivity experiment. In each curve with connecting lines between the data points, the x-axis shows the center of the time bin in which saccades occurred relative to stimulus onset (see Fig. 11A), and the y-axis shows our measure of the drift response strength (Materials and Methods). The floating data points (and associated horizontal dashed lines) in each plot show the no-saccade drift response strength for a given condition (e.g. gray curves in Fig. 11). Each color shows one tested contrast, and error bars denote SEM. As can be seen, the drift response magnitude was suppressed for saccades occurring near stimulus onset and recovered for farther saccades (n >= 97, 149, 105 trials in monkeys A, F, and M, respectively, across all conditions of the experiment). **(D-F)** Similar results for the size tuning experiment (n >= 112, 182, or 50 trials across all conditions in monkeys A, F, and M, respectively).

Therefore, evoked visual responses mediating the drift response are likely suppressed by the presence of peri-stimulus saccades, much like visual responses in some oculomotor areas including the SC (Hafed and Krauzlis, 2010; Chen and Hafed, 2017; Fracasso et al., 2023). Of course, we are not suggesting at all that SC responses mediate the drift response, especially given the results of Fig. 4. Rather, our results mean, instead, that other visual responses impacting the oculomotor system must exhibit saccadic suppression, and it would be interesting to identify in the near future which of these visual responses mediate the drift response.

## Discussion

Ocular position drift eye movements have interested and intrigued neuroscientists for many decades (Ratliff and Riggs, 1950; Barlow, 1952; Nachmias, 1959, 1961; Kowler and Steinman, 1979a, b). The interactions between these eye movements and exogenous sensory events have, however, garnered significantly less attention. We recently observed a robust stimulus-driven ocular position drift response for some visual stimuli (Malevich et al., 2020), and our goal in the present study was to investigate its functional properties much more deeply. Such investigation provides an important foundation for pinpointing the neurophysiological mechanisms giving rise to this drift response, which is itself an important endeavor given how little knowledge we currently have about the neural control of ocular position drifts in general.

Our investigation revealed several interesting properties of the drift response, most notable of which is its robustness even for small foveal and peripheral visual stimuli. There was always a subtle, predominantly upward deviation in ocular position drift trajectories with such stimuli. Given that this deviation alters the spatio-temporal patterns of images impinging on the retina (Kuang et al., 2012; Rucci and Victor, 2015; Ahissar et al., 2016), this suggests that visual onsets in a variety of neuroscientific and cognitive experiments can have sensory representational changes embedded within them, which are directly mediated by stimulus-driven ocular position drifts (in addition to whatever other experimental variables that were being considered by the experimenters). This idea has an interesting parallel in the field of microsaccades; in that related field, it has been suggested that these tiny eye movements can have a significant impact on interpreting various perceptual and cognitive phenomena (Hafed, 2013; Chen et al., 2015; Hafed et al., 2015; Tian et al., 2016).

The ubiquitous nature of the upward velocity pulse that we observed under a variety of conditions might suggest that it is a reflexive eye movement. However, it seems to be too small to be related to a potential dorsal light reflex in lower animals (Brodsky, 1999), and it is also binocular (Malevich et al., 2020) and occurring under binocular visual stimulation conditions. The drift response is also not a general gaze position response to darkness (Malevich et al., 2020). Nonetheless, in the same general theme of linking ancient reflexes to effects in primate vision (Brodsky, 1999), the drift response might help us to learn about low-level, evolutionarily old components of the oculomotor control network, which are still present and active in the primate brain. In fact, given the discrepancy between the results of Fig. 4 and our original hypothesis about the SC mediating the drift response (Malevich et al., 2020), we now seriously ponder the possibility that visual responses downstream of the SC might be more important for observing this response. This might explain why the drift response happens so ubiquitously across many different stimulus types, since visual responses downstream of the SC are bound to influence eye movements, if ever so subtly (by mere proximity to the final oculomotor muscle drive).

Having said that, the drift response as we defined it in the introduction (Fig. 1) is not the only ocular position drift phenomenon that takes place after the onset of small, localized visual stimuli. Indeed, our results from Figs. 4-6 clearly show that there can be spatially-directed drift modulations reflecting the location of a peripheral visual stimulus. This is consistent with our earlier observations about ocular position drifts in peripheral Posner-like cueing tasks (Tian et al., 2018). An important implication of this is that ocular position drifts are not entirely random movements, consistent with other evidence (Murphy et al., 1975; Kowler and Steinman, 1979b, a; Ahissar et al., 2016; Tian et al., 2018; Skinner et al., 2019; Bowers et al., 2021; Reiniger et al., 2021; Clark et al., 2022; Nghiem et al., 2022). This evidence again has parallels in the field of microsaccades, which were thought to be random until two decades ago (Hafed and Clark, 2002; Engbert and Kliegl, 2003).

Mechanistically, spatially-directed drift modulations can emerge from readout of topographically organized visual-motor maps, like in the SC (Robinson, 1972; Ottes et al., 1986; Chen et al., 2019). For example, we recently found that at the time of saccade triggering, even spontaneous spiking in movement-unrelated locations of the SC map can be instantaneously readout by the oculomotor system to modify the flight trajectory of saccades (Buonocore et al., 2021). In a similar light, spatial readout of the entire landscape of SC activity can dictate the smooth position deviations during gaze fixation, and such landscape will have clear spatial biases when some SC neurons discharge visual bursts after localized, peripheral stimulus onsets. The spatially-directed drift effects that we observed would then reflect these biases. Such a mechanism would be consistent with how the SC contributes to the much faster (relative to the drift response) smooth pursuit eye movements in general, like when tracking an invisible moving goal that is being represented in a spatially broad manner across the SC map (Hafed and Krauzlis, 2008). Such a mechanism would also be consistent with the idea that the upward drift pulse that accompanies spatially-directed drift modulations can be mediated by some other circuit operations (potentially even downstream of the SC).

Returning to the more reflex-like, predominantly upward drift response (Fig. 1), as we said, it is likely dissociated from SC activity because it remains predominantly upward even when SC neurons representing the lower visual field are expected to be bursting after stimulus onset (Fig. 4). This idea can and should be explicitly tested by recording SC activity from the same task of Fig. 4. We also think that other evidence in our data could point to a dissociation of the drift response from SC activity. Specifically, we often observed a transient eye position modulation right before the upward velocity pulse, a clear example of which is seen in Fig. 1B, C. Such a transient modulation jumpstarts the whole drift response sequence, and it seems to also be feature-tuned. That is, it was modulated in strength and timing as a function of some stimulus properties, like size and contrast (Figs. 2, 3). This could suggest that visual bursts mediating the drift response (wherever they may actually be in the end) could initially cause such transients, and that the subsequent upward drift pulse could reflect various time constants of the oculomotor control network and oculomotor plant (Robinson, 1964). For example, using a systems control perspective, imagine a negative feedback control loop driving an eye plant, and now drive the whole circuit with a temporal impulse function. Part of the resulting response would reflect the time constants of not only the control loop but also the eye plant. If that is the case, then future experiments need to understand why driving the oculomotor control network with a temporal impulse function (a brief visual burst) would eventually lead to a predominantly upward eye movement, as opposed to downward or horizontal or in some random direction, after the initial transient modulation.

Regardless of the mechanism, all of the above evidence suggests that the drift response falls in a class of eye movement phenomena that may be evoked directly by visual bursts in the oculomotor system, as we recently discussed (Buonocore and Hafed, 2023; Khademi et al., 2023). These phenomena also include express saccades (Fischer and Boch, 1983; Edelman and Keller, 1996; Marino et al., 2015; Hall and Colby, 2016) and saccadic inhibition (Reingold and Stampe, 1999, 2002, 2004; Edelman and Xu, 2009; Khademi et al., 2023). In fact, we think that saccadic inhibition and the drift response are likely mediated by the same structures (Figs. 7-9), further emphasizing the idea that the drift response might be reflexive. If so, one might make some neurophysiological predictions here. Specifically, if the hypothesis (Hafed et al., 2021b; Buonocore and Hafed, 2023) holds that omnipause neurons in the brainstem have visual pattern responses explaining the feature tuning properties of saccadic inhibition, and if drift responses are also triggered by these neural bursts, then one prediction is that visual bursts in these omnipause neurons might act as the “temporal impulse function” that jumpstarts the drift response, which we alluded to above. If so, this would implicate omnipause neurons in more than just the interruption of saccades (Keller and Edelman, 1994; Kaneko, 1996; Keller et al., 1996; Gandhi and Keller, 1999), and the next question will be why brief burst impulses in omnipause neuron activity could cause a small, but smooth, eye position deviations (in addition to inhibiting saccade generation).

Finally, regardless of whether these ideas are experimentally validated or not, it is also important to consider our observation that the drift response was suppressed by the occurrence of peri-stimulus saccades (Figs. 11, 12). Some smooth eye movement phenomena are actually enhanced when stimuli occur right after microsaccades (Chen and Hafed, 2013), but these phenomena typically involve ocular following of moving stimuli (Chen and Hafed, 2013). In our case, the drift response was not to follow a moving target or pattern. Its suppression, thus, predicts that visual bursts mediating the drift response (wherever they may be) must be suppressed by peri-stimulus saccades. It would be interesting to also test for this idea neurophysiologically.

### Grants

We were funded by the following grants from the Deutsche Forschungsgemeinschaft (DFG; German Research Foundation): BU4031/1-1, HA6749/4-1, BO5681/1-1, HA6749/3-1, and SFB 1233, Robust Vision: Inference Principles and Neural Mechanisms, TP11, project number: 276693517.

